# Reconfiguration of genome-lamina interactions marks the commissioning of limb cell-fates

**DOI:** 10.1101/2025.05.05.651267

**Authors:** Konrad Chudzik, Isabel Guerreiro, Samy Kefalopoulou, Alex Abraham, Magdalena Schindler, Alessa Ringel, Mario Nicodemi, Irina Solovei, Andrea M. Chiariello, Stefan Mundlos, Jop Kind, Michael I. Robson

**Affiliations:** Max Delbrück Center for Molecular Medicine, Berlin Institute for Medical Systems Biology, Berlin, Germany; Max Planck Institute for Molecular Genetics, Berlin, Germany; Hubrecht Institute, Royal Netherlands Academy of Arts and Sciences (KNAW) and University Medical Center Utrecht, Utrecht, the Netherlands; Oncode Institute, The Netherlands; Dipartimento di Fisica, Università di Napoli Federico II and INFN Napoli, Complesso Universitario di Monte Sant’Angelo, Naples, Italy; Department of Biology II, Biozentrum, Ludwig-Maximilians University Munich (LMU), Planegg-Martinsried, Germany; Department of Molecular Biology, Radboud Institute for Molecular Life Sciences, Radboud University Nijmegen, Nijmegen, the Netherlands

## Abstract

Diverse forms of heterochromatin block inappropriate transcription and safeguard differentiation and cell identity. Yet, how and when heterochromatin is reconfigured to facilitate changes in cell-fate remains a key open question. Here, we address this by mapping a prevalent heterochromatic feature - genome-lamina interactions - relative to transcription in single-cells during mouse embryogenesis. We find that genome-lamina interactions remain relatively uniform between germ layers following gastrulation but are extensively reconfigured in diverse tissues during later organogenesis. Focusing on limb development, we demonstrate that genome-lamina interactions are selectively released in early multipotent progenitors at key developmental genes and their surrounding regulatory domains. This “lamina-release” often precedes gene expression at later developmental stages, suggesting it primes regulatory domains for future potential activation. Conversely, lamina-release coincides with chromatin opening at sites of crucial limb transcription factor binding, and so is closely intertwined with the regulatory machinery driving limb formation. Finally, we show that the boundaries of topologically-associated domains (TADs) constrain the spread of lamina-release at a limb gene locus. This ensures independent lamina dynamics between neighbouring domains. Together, our data suggest a previously unrecognised process where genome-lamina interactions are selectively released at regulatory domains to transition loci toward more permissive chromatin states, thereby potentiating cell type specific activation. Our work thus reveals how systematic heterochromatin reorganization links to developmental multipotency, providing mechanistic insights into cell-fate decisions *in vivo*.

## MAIN

During embryogenesis, precise changes in gene expression are essential for determining cell identity and the correct development of tissues and organs^1^. Limb development exemplifies this principle, as distinct spatiotemporal gene patterns drive progenitor cells to differentiate into diverse cell types, culminating in a highly structured limb anatomy^2,3^. By definition, such precision requires gene repression, which is achieved by loci adopting diverse heterochromatic states that block TF binding and inappropriate transcription^4–7^. However, this raises a key unanswered question - what events transition loci between heterochromatic and euchromatic states to facilitate changes in activity and cell-fate?

Studies addressing this have largely focussed on examining how heterochromatin is remodelled at the level of histone modifications to regulate genes and differentiation^4,7^. Yet, there is an equally fundamental aspect of heterochromatin whose developmental dynamics remain elusive - the spatial positioning of genes in the nucleus. Many inactive loci are positioned at the nuclear periphery by interacting with the nuclear lamina in large lamina-associated domains (LADs)^8^. LADs are typically enriched in a compacted, H3K9me2-marked type of heterochromatin that contains thousands of genes held in a generally repressed state^8–10^. Accordingly, LADs display considerable cell type specificity in several *in vitro* differentiation models, with many loci losing genome-lamina interactions when active and gaining them when repressed^11–16^. This also appears functionally significant as disrupting LADs impairs gene regulation, differentiation, and cell identity in the largely *in vitro* contexts tested so far^11,13,16–19^.

Despite this, the function of LAD reorganization in gene regulation and development is yet to be determined. First, we do not know when LADs are reorganized during differentiation, nor whether this remodelling drives or passively follows altered gene activities^11,13,16–18,20–25^. Second, it is unresolved how LAD dynamics relate to the wider chromatin machinery that governs gene expression. Developmental genes are regulated by transcription factors (TFs) binding to enhancers found in extended gene regulatory landscapes that can span hundreds or thousands of kilobases^26,27^. These landscapes are spatially confined within 3D topologically-associating domains (TADs) that favor enhancer interactions with correct target genes^27,28^. Yet, how LAD reorganisation unfolds relative to these highly interconnected features of regulatory landscapes is poorly explored, particularly in native *in vivo* contexts^29,30^. Deciphering how LADs progressively reconfigure is thus vital, both for understanding how genomes function but also how cells control their fates.

### LADs extensively reconfigure during organogenesis

Addressing this challenge requires measuring single-cell LAD and transcriptional dynamics along multiple developmental trajectories *in vivo*. We consequently implemented scDam&T-seq^31^ in mice to simultaneously map LADs and transcription together in single cells from complex tissues during mouse embryonic development (Fig. 1a and ED Fig. 1a-b). To this end, we created a transgenic mouse line that conditionally expresses the DNA adenine methyltransferase (Dam) fused to a component of the nuclear lamina (LMNB1) (see Methods). Once expressed, Dam-LMNB1 labels LADs in cells which can then be individually isolated from embryos by fluorescence activated cell sorting (FACS) for scDam&T-seq processing (ED Fig. 1a-b). Critically, no Dam-LMNB1 activity was detected in the absence of tamoxifen (ED Fig. 1c), while tamoxifen injections resulted in strong induction in multiple tissues and across embryonic stages (ED Fig. 1c-e). An injection time course also demonstrated that 92.5% of cells induce Dam-LMNB1 construct expression between 8-17 hours after tamoxifen delivery (ED Fig. 1c). This provides an ∼9 hour time window where Dam- LMNB1 labels LADs, enabling their dynamics to be profiled with high temporal resolution during embryogenesis *in vivo*.

**Figure 1.**
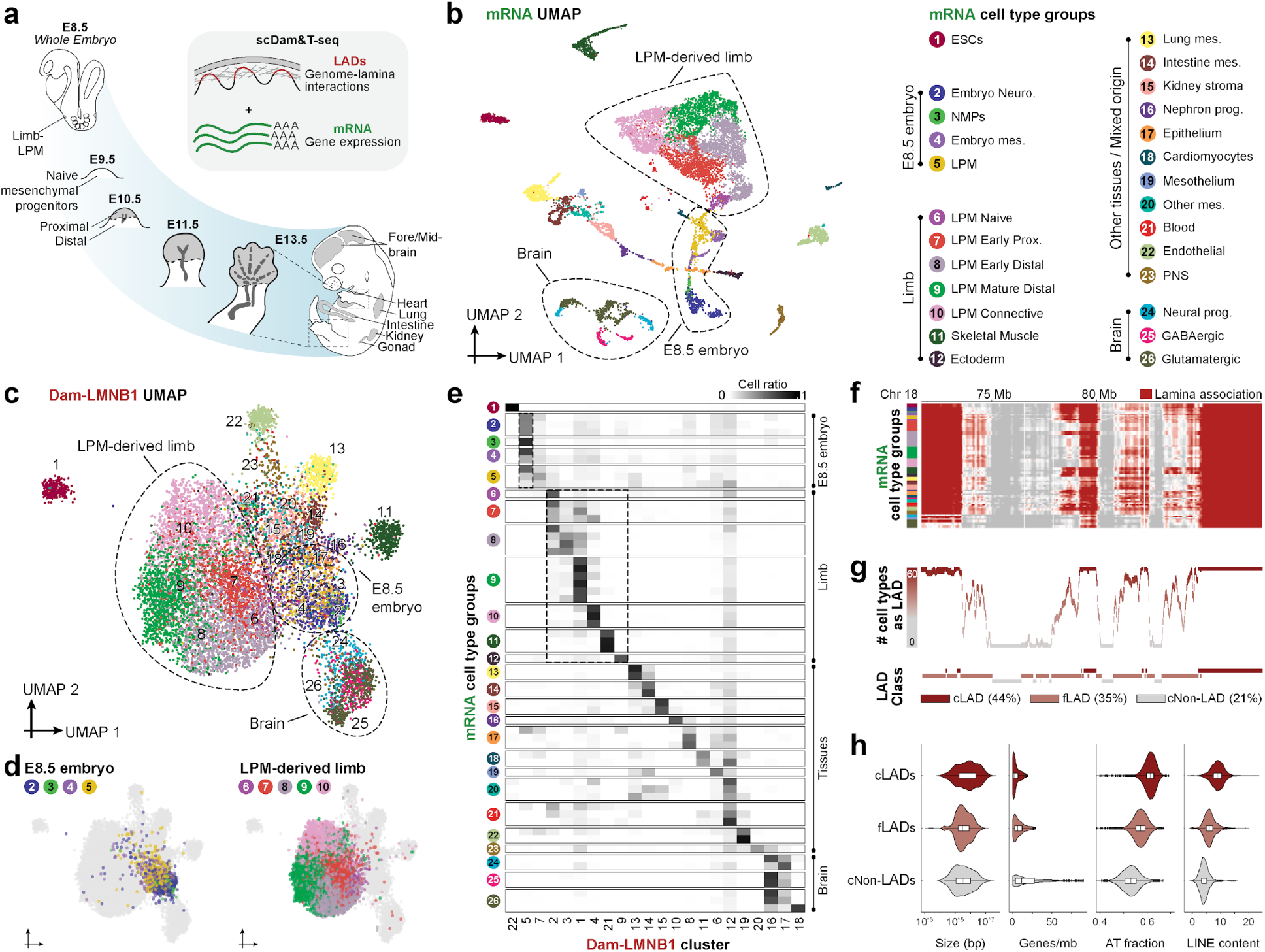
LADs reconfigure at developmental genes during organogenesis. a, Schematic of mouse limb development and tissues sampled with scDam&T-seq. **b-d,** scDam&T-seq uniform manifold approximation and projection (UMAP) visualization of single cells from sample tissues. Coordinates are defined by mRNA (n = 11330) (b) or Dam-LMNB1 signal (n = 10974) (c,d). Colours and numbers in both UMAPs correspond to mRNA cell type groups. Neuro., neuroectoderm. Mes., mesenchyme. LPM, lateral plate mesoderm. Prox., proximal. Prog., progenitors. **e,** Heatmap of correspondence between mRNA-defined cell types and Dam-LMNB1 cell clusters. mRNA cell types are organised into 26 cell type groups for clarity. **f,** Tracks of binarised lamina association in a representative genomic region for each mRNA cell type with their broader cell type groups highlighted. **g**, Matching classification of genomic regions (x axis) into cLADs, fLADs and ciLADs based on their frequency of interaction with the lamina across cell types (y axis). **h,** Quantification of sequence properties of LAD classes (Size: cLADs, n = 627, fLADs, n = 1276; cNon-LADs, n = 568; Genes/mB: cLADs n = 374, fLADs, n = 837; cNon-LADs, n = 406; AT fraction: cLADs, n = 0.518; fLADs, n = 0.403; cNon-LADs, n = 0.248; LINE content: cLADs, n = 627; fLADs, n = 1276; cNon-LADs = 568). See also ED Fig. 1-3 and Tables 1-4.

After establishing experimental conditions, we next applied scDam&T-seq to mouse embryogenesis, obtaining 3000 cells from multiple E13.5 tissues where initial organ formation has already occurred (Fig. 1a). These include lung (583), heart (107), kidney (446), intestine (668), gonads (218), forebrain (364) and midbrain (614), as well as 314 embryonic stem cells (ESC) for comparison. We also particularly focused on the developing limb where precise spatiotemporal changes in gene expression drive progenitors along multiple developmental lineages (Fig. 1a)^2,3^. Specifically, the lateral plate mesoderm (LPM) generates bone, cartilage, and connective tissue while other migrating progenitors produce skeletal muscle, neurons, and endothelial cells^2,3^. In total, this yielded 11330 cells which passed thresholds for robust joint mRNA and LAD profiling (ED Fig. 1f-i). Processed scDam&T-seq samples and quality statistics are listed in Tables 1-2.

First, we examined differences in LAD patterns between all cell types and tissues. We used the single-cell mRNA modality to cluster and annotate cells based on marker genes and published atlases (see Tables 3-4 and Methods). This identified 60 cell types, which we classified into 26 broader cell type groups based on their developmental origins (Fig. 1b and ED Fig. 2a,b). To assess LAD differences across single cells, we also separately clustered the same cells by their Dam-LMNB1 signal which identified 22 distinct Dam-LMNB1 cell clusters (ED Fig. 2c). Quantifying the overlap between mRNA cell type groups and Dam-LMNB1 cell clusters revealed that LADs distinguish cells by tissue, developmental stage, and cell-identity to different extents (Fig. 1c-e and ED Fig. 2d-e). In particular, all four cell type groups identified in the E8.5 embryo intermingle in a single Dam-LMNB1 cluster, cluster 5 (Fig. 1d,e and ED Fig. 2c). This shows that LAD patterns of diverse cell types at E8.5 are surprisingly similar. In contrast, we observe that LADs diverge at later stages with cells from multiple tissues forming distinct Dam-LMNB1 clusters (Fig. 1c-e and ED Fig. 2c-e). For example, limb cells distribute into six distinct Dam-LMNB1 clusters that correspond to major limb cell type groups. This includes LPM-derived naive progenitors, which segregate from the more differentiated cells they give rise to (Fig. 1d,e). Combined, this supports that LADs are unexpectedly similar in cells from fundamentally different germ layers following gastrulation. Yet, during later organogenesis, LADs reconfigure resulting in cell type-specific genome-lamina interactions.

**Figure 2.**
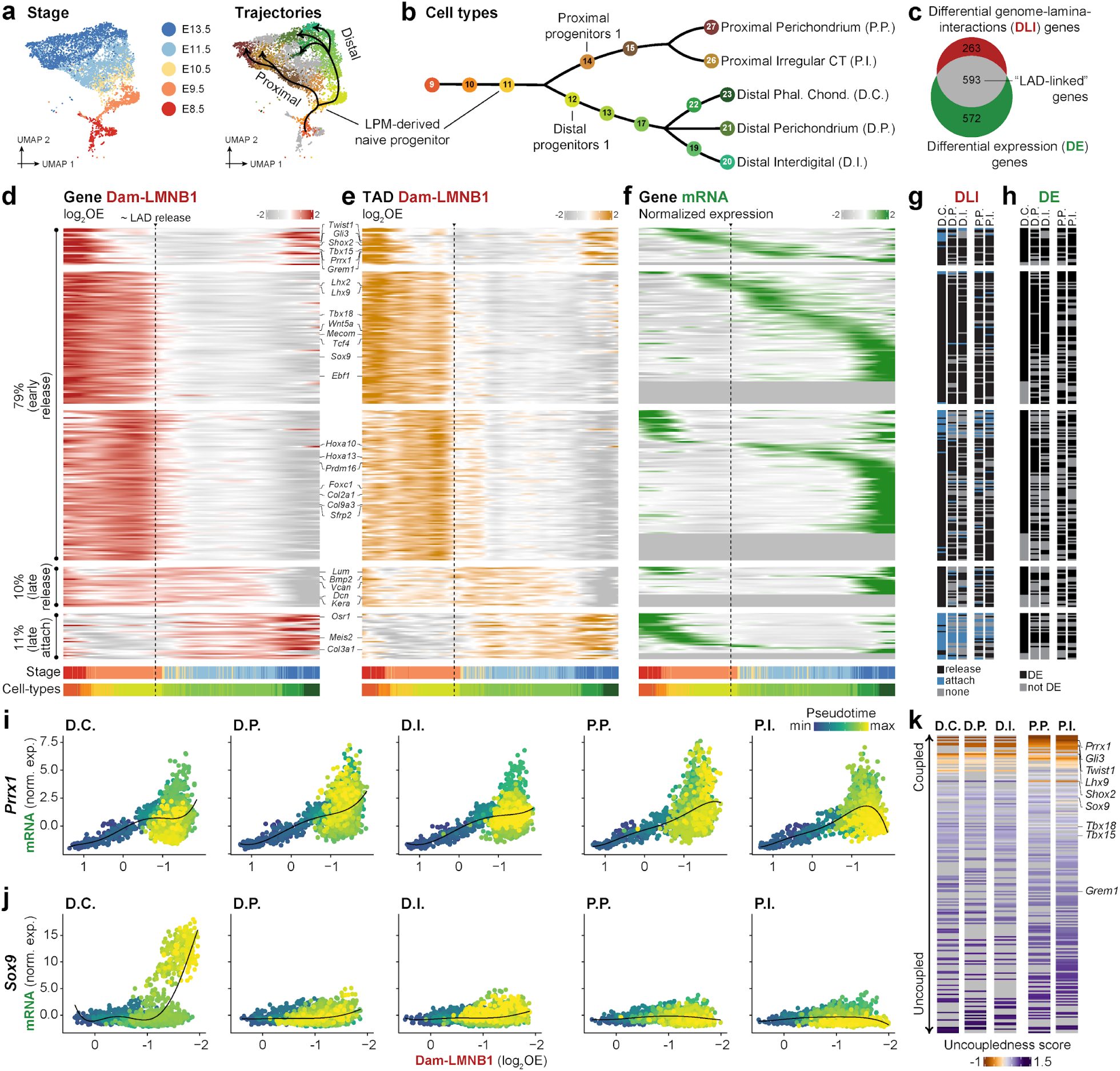
Genes and their regulatory domains release the lamina prior to activation. a, UMAP visualisations of LPM-derived limb cells with coordinates defined by mRNA and cells coloured by embryonic stage (left) or cell types of selected trajectories (right) (n = 6486). Arrows indicate paths of selected proximal and distal differentiation trajectories toward E13.5 limb cell types. **b,** Schematic dendrogram of selected trajectories that arise from a common multipotent naive limb progenitor. Numbers refer to annotated cell types, which are fully listed in ED Fig. 2. **c,** Venn diagram of genes with differential genome-lamina-interactions (DLI) or differential gene expression (DE) in at least one trajectory. **d-f,** Heatmap showing LAD-linked genes (y axis) and their differential Dam-LMNB1 (d, e) or mRNA signals (f) along pseudotime (x axis) in the D.C. trajectory. Dam-LMNB1 signal is plotted for the 200kb bin surrounding each gene (d) or their surrounding TADs (e). Genes are shown if they exhibit DLI and DE in at least one trajectory. Example genes with known limb functions are highlighted. **g-h,** Corresponding classification of LAD-linked genes as DLI (g) and/or DE (h) in indicated trajectories. **i-j.** Relationship between Dam-LMNB1 and mRNA dynamics for *Prrx1* (i) and *Sox9* (j) across indicated trajectories. **k,** Uncoupledness scores of lamina and transcriptional dynamics for LAD-linked genes across trajectories. See also ED Fig. 4-6 and Tables 5-6.

We then further characterised the extent of LAD variability across the sampled tissues by comparing average pseudobulk LAD profiles for all cell types (Fig. 1f). This revealed that 21% of the genome represents regions that never contact the lamina, which are termed constitutive non-LADs (cNon-LADs) while 44% represents regions with constitutive genome-lamina interactions (cLADs) (Fig. 1g). The remaining 35% of the genome are facultative LADs (fLADs) that are gained or lost in at least one cell type and show cell type specificity. Compared to ciLADs, fLADs are more gene poor, AT-rich, and enriched in LINEs and this difference is even more pronounced in cLADs (Fig. 1h). The cell type specificity of LAD patterns is thus associated with specific genomic signatures, as reported previously in *in vitro* studies^9,32^.

We then sought to link differences in LAD patterns to specific cell types and to identify the genes these genomic regions contain. For this, we clustered genomic bins according to Dam-LMNB1 signal variation across all single cells (ED Fig. 3a). This identified 16 classes of genomic regions with distinct lamina behaviours (ED Fig. 3b,c). For example, class 2 regions represent cLADs while class 13 regions are fLADs that contact the nuclear lamina only in ESCs (ED Fig. 3d). Strikingly, different classes contain genes with gene ontology (GO) terms related to their cell type-specific LAD behaviours (ED Fig. 3e). Class 4 LAD regions undergo lamina-release specifically in limb LPM-derived cell types and are enriched in genes related to limb morphogenesis. Likewise, class 11 LAD regions selectively release from the lamina in brain cell types and contain genes linked to axon guidance and neuron fate commitment (ED Fig. 3e). In short, much of the genome shows tissue-specific genome-lamina interactions during organogenesis and this is linked with relevant developmental genes.

**Figure 3.**
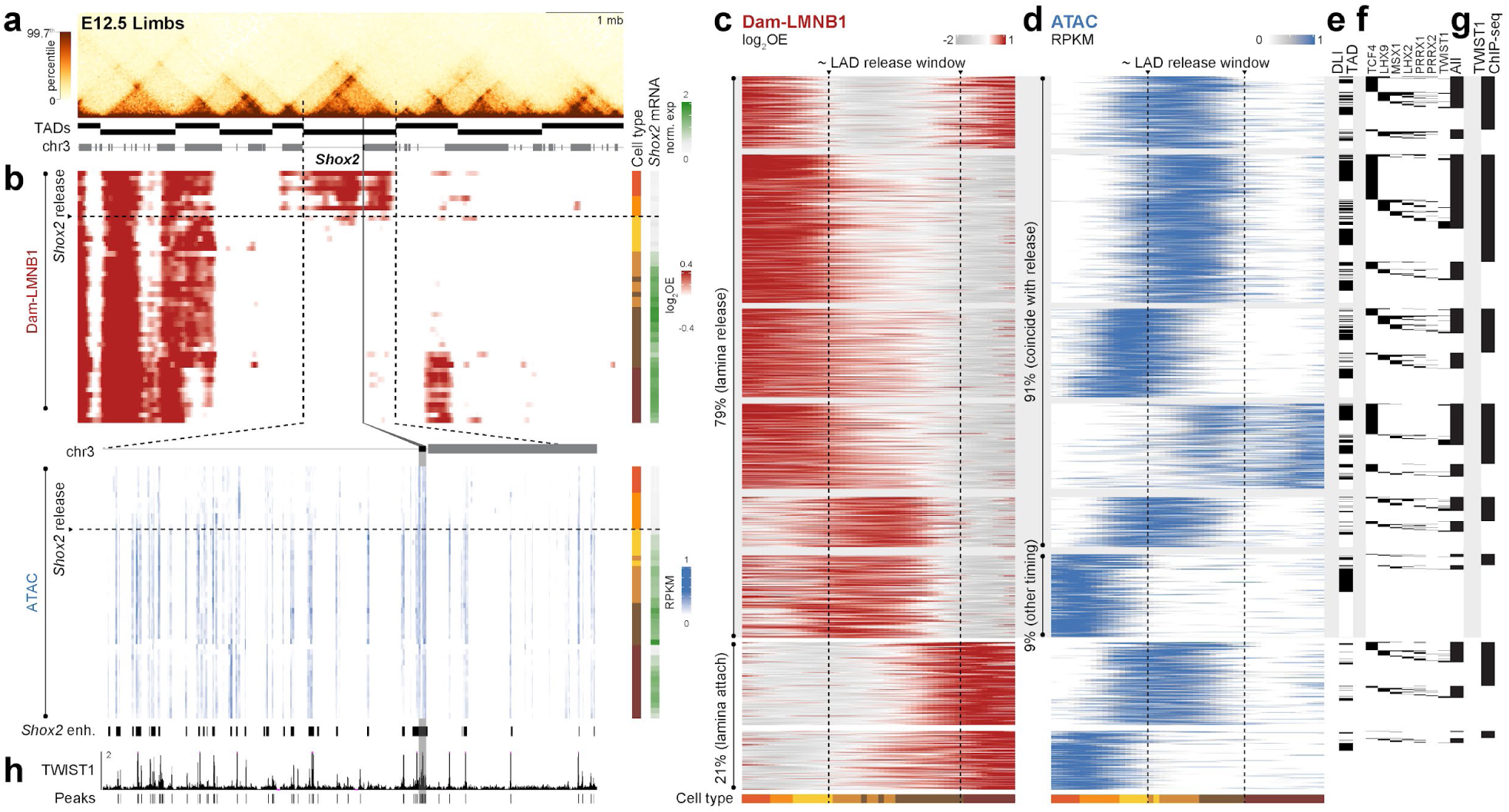
Lamina-release coincides with chromatin accessibility and limb TF binding. **a**, E12.5 limb Hi-C^44^ of the *Shox2* locus with called TADs (top) and gene track (bottom). **b**, Corresponding Dam-LMNB1 and ATAC signal (x axis) in single cells ordered along pseudotime (y axis) in the P.I. trajectory. *Shox2* mRNA signal per single cell is also shown (rightmost barplot). *Shox2* and its TAD are highlighted by the vertical gray bar and dotted lines, respectively. Lower single-cell ATAC heatmap is zoomed to the *Shox2* locus, with characterised *Shox2* enhancers shown below^46,47^. **c-d,** Heatmap of changing Dam-LMNB1 (c) and ATAC (d) signal for genomic bins that show both accessibility and lamina interaction dynamics (y axis) along P.I. trajectory pseudotime (x axis). Cell types are indicated below. **e-g,** Corresponding overlap of ATAC peaks with TADs of LAD-linked genes (e), predicted TF binding (f), or TWIST1 ChIP-seq peaks in E10.5 limbs^45^ (g). **h,** TWIST1 E10.5 forelimb ChIP-seq at the *Shox2* locus with called peaks below^45^. See also ED Fig. 7-10 and Tables 7-10.

### Gene regulatory domains undergo lamina-release in progenitors that precedes later transcriptional activation

We next clarified the timing of LAD reconfiguration and its relationship to gene expression and cell-fate by focussing on the developing limb. In the limb, scDam&T-seq captured the early specification of LPM-derived naive progenitors (E8.5-E9.5) and their bifurcation into proximal and distal precursors (E9.5-10.5) (Fig. 2a and ED Fig. 4a)^2,3,33^. From E10.5 to E13.5, these precursors further differentiate into proximal and distal cell types along increasingly branching trajectories. We identified five of these differentiation trajectories where successive cell types could be confidently verified based on previous literature (Fig. 2b and ED Fig. 4b) (see Methods). These include two proximal trajectories, for irregular connective tissue (P.I.) and proximal perichondrium (P.P.), and three distal trajectories, for phalange chondrocytes (D.C.), interdigital mesenchyme (D.I.), and distal perichondrium (D.P.)^33^. Using the transcription modality of scDam&T-seq, we then ordered cells from each of these trajectories along pseudotime (ED Fig. 4c). In this way, the selected limb trajectories represent multiple lineages and branch points to contrast LAD and transcriptional dynamics over developmental time.

**Figure 4.**
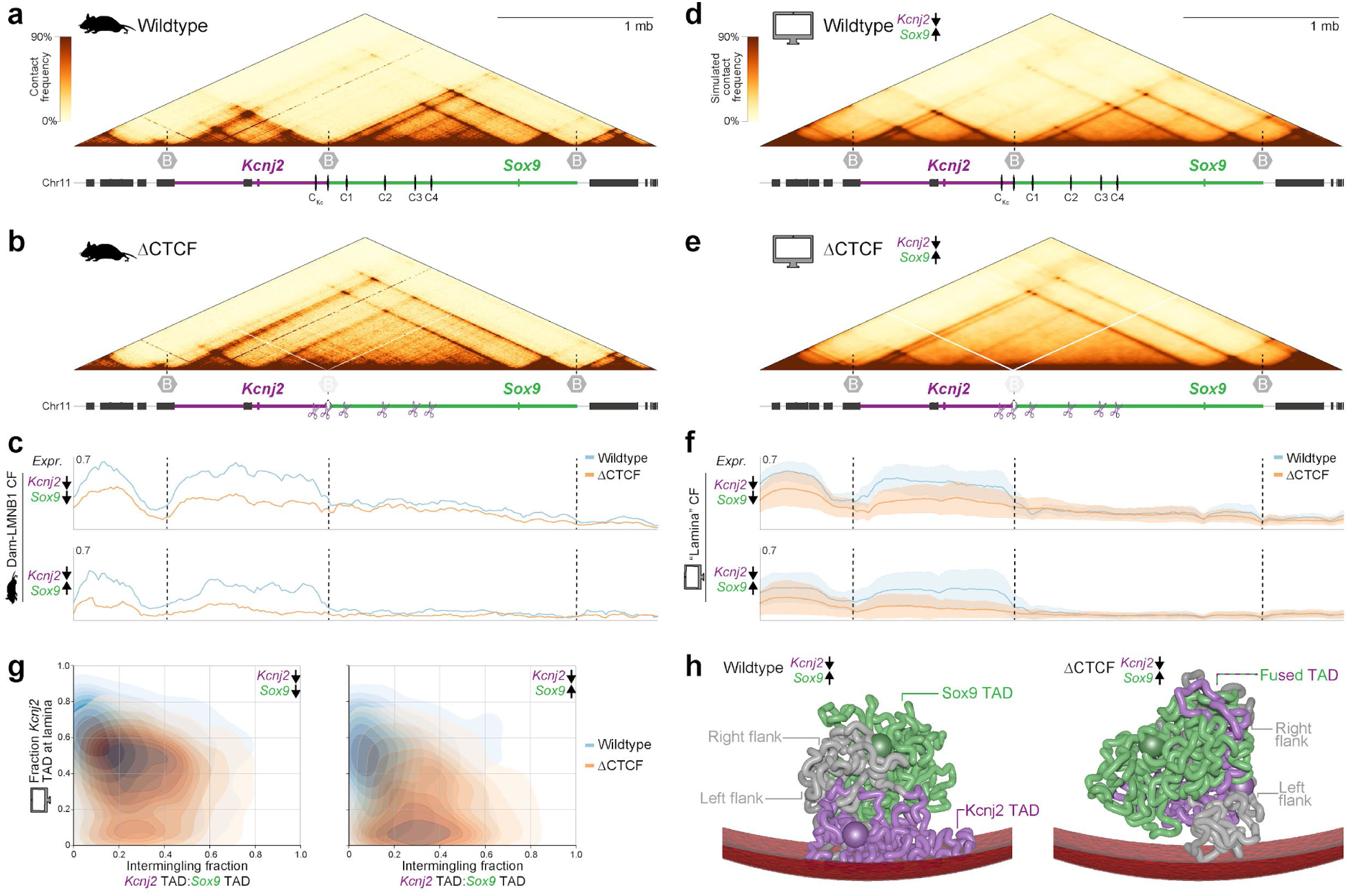
**TAD CTCF boundaries enable independent LAD dynamics by blocking chromatin intermingling. a-b**, cHi-C of the *Kcnj2/Sox9* locus in wild-type (A) or ΔCTCF (B) E12.5 limbs with genes and CTCF sites highlighted below^50^. **c**, Average Dam-LMNB1 contact frequency (CF) in wildtype and mutant E13.5 LPM-derived limb cells grouped by low (↓) or high (↑) *Kcnj2*/*Sox9* expression level. CF reflects the fraction of cells for which a genomic bin contacts the lamina. **d-f**, Simulated Hi-C (D-E) and lamina CF (F) in wildtype or ΔCTCF mutant limb cells. **g**, Matching model quantification of the simulated *Kcnj2* region’s lamina contact and intermingling with the *Sox9* domain. n = 8680 simulated structures per condition. **h,** Representative polymer models of locus in wildtype or ΔCTCF mutant limb cells with low *Kcnj2* and high *Sox9* expression. See also ED Fig. 11, Videos 1-2, and Table 12.

Using the variation in Dam-LMNB1 signal over pseudotime, we first globally identified genes with differential genome-lamina interactions (DLI) across any of the 5 trajectories. This revealed 856 genes with DLI in at least one trajectory (Fig. 2c). Of these, 593 (71%) are also differentially expressed (DE). In total, roughly half (51%) of the 1165 genes that show differential expression across any of our limb trajectories also display dynamic genome-lamina interactions. We hereafter refer to these genes as “LAD-linked” genes, which are listed in full in Table 5.

We then evaluated the Dam-LMNB1 signal for LAD-linked genes over pseudotime in each trajectory (Fig. 2d and ED Fig. 5a). This revealed a strikingly uniform wave of lamina-release at the origin of all five trajectories where distal and proximal lineages diverge. Specifically, a majority (58-81%) of LAD-linked genes begin releasing from the lamina in naive progenitors and are completely released by the first distal or proximal precursors (Fig. 2d and ED Fig. 5a). Notably, early lamina-release genes include key TFs and signalling factors that are essential for limb development, such as *Twist1, Gli3, Shox2, Tbx15, Prrx1, Grem1, Lhx2, Lhx9, Mecom, Sox9, Runx2, Tbx18*^34–43^. By contrast, only a small minority of remaining LAD-linked genes show alternative dynamics, including 7-11% that attach to the lamina only in later cell types (Fig. 2d and ED Fig. 5a). Thus, LADs at key limb genes are selectively released in a synchronous wave in early limb progenitors. We also note that 49% of these releasing genes detach the lamina in both distal and proximal precursors (Fig. 2g and ED Fig. 5b). This shows that, despite diverging, both proximal and distal lineages release a substantially overlapping set of genomic loci.

We additionally tested whether lamina-release involves the local detachment of the gene itself or occurs in the context of a broader genomic region. In particular, we focussed on whether LAD release extends across surrounding TADs that frequently constitute the regulatory domains of developmental genes^26,27^. Using published E12.5 forelimb Hi-C data^44^, we identified TAD boundaries and displayed changes in the Dam-LMNB1 signal for LAD-linked genes and their TADs over pseudotime. TADs displayed the same timing of lamina-release as their hosted LAD-linked genes in both proximal and distal lineages (Fig. 2e and ED Fig. 5a). This broad lamina-release also appears to be constrained to the TADs themselves. Indeed, the strongest loss of genome-lamina interactions is observed within TAD boundaries when LAD-linked genes are released from the lamina (ED Fig. 5c). Collectively, these data shows that genes release the lamina together with the TAD regulatory landscapes that, in many cases, control their expression.

We finally assessed how lamina-release corresponds with gene expression dynamics over pseudotime. As expected of LADs repressive nature^9^, a subset of genes activate transcription simultaneously with lamina-release in distal and proximal progenitors (Fig. 2d,f,i and ED Fig. 5a). This includes the key limb TF genes *Prrx1*, *Twist1, Lhx9, and Tcf4*^34,38,40,45^, suggesting that TFs driving limb differentiation are the first to activate following lamina-release.

However, many LAD-linked genes exhibited striking delays between lamina-release and transcriptional activity, while others remained inactive in specific trajectories despite detaching the lamina (Fig. 2d-h and ED Fig. 5a). For example, *Sox9* lamina-release occurs in the early progenitors of all trajectories (Fig.2j and ED Fig. 4c). Yet, it only strongly activates four days later in phalange chondrocytes in the DC trajectory, while remaining inactive in all others. To quantify this delay, we developed an "uncoupledness score" to globally measure the temporal disparity between LAD dynamics and transcription for each gene (ED Fig. 6, see Methods). This revealed that lamina-release and transcription are indeed temporally uncoupled at most LAD-linked genes (Fig. 2k). LAD release in early limb progenitors thus often precedes the potential for a gene’s later transcriptional activity.

In summary, the scDam&T-seq time series reveals that LADs are extensively reorganized in early limb progenitors. This reorganization is targeted to key developmental genes and their regulatory domains, the majority of which detach from the lamina prior to their potential activation in later cell types. Combined, this suggests that tissue-specific LAD reconfiguration in progenitors primes loci for future activity, thereby commissioning new cell-fate decisions for proper spatiotemporal limb development.

### Lamina-release is linked to the binding of key early limb TFs

TADs are often populated by multiple regulatory elements that activate developmental genes^26,27^. We consequently hypothesised that the observed lamina-release is connected to TF binding at regulatory elements distributed throughout detaching TADs, as proposed previously for specific loci *in vitro*^30^. To test this, we generated single-nucleus chromatin accessibility and mRNA 10x Multiome datasets of the same time points as collected previously for scDam&T-seq. This includes whole E8.5 embryos and E9.5, 10.5, 11.5 and 13.5 limbs. This yielded a total of 14817 10x Multiome nuclei which were clustered into 54 cell types based on the mRNA modality (see Methods and Tables 7-10). 38 equivalent cell types independently annotated in the 10x Multiome or scDam&T-seq datasets clustered together when integrated by their common transcriptional readout (ED Fig. 7 and Tables 10-11). The integrated dataset allowed us to reconstruct representative proximal (P.I.) and distal (D.C.) limb trajectories, providing a shared pseudotime axis for both Dam-LMNB1 and ATAC readouts (ED Fig. 8a,b). This approach enables a direct comparison of changes in genome-lamina interactions, accessibility, and transcription during limb development.

To evaluate the dynamics of the three modalities over pseudotime, we first focussed on a specific locus of a LAD-linked gene, *Shox2*. This revealed a striking correlation between lamina-release and increasing chromatin accessibility in both trajectories. As naive progenitors transition into proximal or distal precursors, *Shox2* and its 1.05 Mb TAD progressively lose Dam-LMNB1 signal (Fig. 3a,b and ED Fig. 9a,b)^44^. Simultaneously, numerous differential ATAC peaks found in *Shox2*’s TAD gain accessibility during the same cell type transitions, many of which overlap previously characterized *Shox2* limb enhancers^46,47^. Combined, this demonstrates that lamina- release at the *Shox2* locus is closely linked to the early opening of regulatory elements throughout its detaching TAD. However, despite this reconfiguration being common to both trajectories, *Shox2* is only active in the P.I. trajectory (Fig. 3b) while it remains silent in the DC lineage (ED Fig. 9b). Lamina-release and regulatory element opening alone therefore do not seem to drive *Shox2* transcription. Instead, our results suggest that these changes establish a chromatin state that is permissive for later potential gene activation.

We further evaluated the genome-wide relationship between increasing chromatin accessibility and lamina-release. To this end, Scenic+ was applied to 10x Multiome nuclei from 7 LPM-derived cell types, up to the proximal and distal precursors where lamina-release is mostly complete (ED Fig. 8c)^48^. Scenic+ identifies eRegulons which are composed of differentially accessible regions, the TF predicted to bind to them, and the genes they are inferred to control (ED Fig. 8c). This analysis identified 133 eRegulons that incorporate 18029 differentially accessible ATAC peaks, of which 2552 display differential lamina interactions (DLI) over pseudotime. As predicted, 72-79% of these DLI ATAC peaks lose Dam-LMNB1 signal in early limb progenitors, coinciding with the major wave of LAD-release observed previously (Fig. 3c and ED Fig. 9c). Matching this, DLI ATAC peaks, which often overlap with previously identified lamina-releasing TADs (Fig. 3e and ED Fig. 9e), frequently gain chromatin accessibility as Dam-LMNB1 signal is lost (Fig. 3d and ED Fig. 9d). Lamina-release is therefore globally tied to the opening of regulatory elements in limb progenitors.

Chromatin accessibility is frequently associated with TF binding. We therefore used our eRegulon analysis to identify TFs predicted to bind to DLI ATAC peaks. This revealed that many DLI ATAC peaks are predicted to be bound by TFs with well-characterised roles in early limb development, such as TCF4, MSX1, LHX2, LHX9, PRRX1, PRRX2, and TWIST1 (Fig. 3f and ED Fig. 9f)^34,40,45,49^.

For the latter, we could directly confirm TWIST1 binding at a majority of differential ATAC peaks using published E10.5 limb ChIP-seq data^45^ (Fig. 3g and ED Fig. 9g). This includes the *Shox2* locus, where TWIST1 binds to multiple characterised *Shox2* enhancers that gain accessibility in *Shox2*’s TAD concurrently with its lamina-release (Fig. 3h and ED Fig. 9h)^46,47^. Thus, early lamina- release is tightly intertwined with the opening of regulatory elements and the binding of key limb TFs throughout regulatory landscapes. This principle even extends to the genes encoding these lamina-release linked TFs. Indeed, the *Lhx2, Lhx9, Prrx1,* and *Twist1* gene regulatory landscapes are all themselves in early detaching LAD regions (Fig. 2d-g) (see ED Fig. 10 for an example).

In sum, this demonstrates that LADs and regulatory element accessibility are extensively reconfigured together across gene regulatory landscapes in early limb progenitors. This reconfiguration precedes the potential for transcriptional activity in future cell-fates, and is connected to the binding of key TFs that drive limb development.

### TAD boundaries partially constrain the extent of lamina-release

Having gained insight into the relationship between regulatory landscapes and LAD release, we finally investigated what prevents this phenomenon from spreading to neighbouring loci. Our previous analysis demonstrated that lamina-release frequently extends up to surrounding TAD boundaries (Fig. 2e and ED Fig. 5c). We therefore hypothesised that TAD boundaries help restrain the extent of lamina-release and so their removal would cause LAD dynamics to spread.

We tested this by exploiting the well-characterised *Kcnj2/Sox9* locus, which we selected for two reasons. First, *Kcnj2* and *Sox9* are located in neighbouring TADs and interact with the lamina independently from one another in limb cell types. *Sox9*’s TAD undergoes LAD-release in early limb progenitors before *Sox9* activates in later phalange chondrocytes (Fig. 4a and ED Fig. 11a,b). By contrast, *Kcnj2*’s TAD largely continues to independently interact with the lamina and remains inactive in the same cells. Second, a previous study created a ΔCTCF mutant mouse line that selectively removed the 11 CTCF sites that separate the *Kcnj2/Sox9* TADs^50^. This causes the TADs to fuse in mutant limbs, thereby ectopically connecting *Kcnj2* with inappropriate *Sox9* enhancers that drive its misexpression (Fig. 4b)^50,51^. Combined, these features create an ideal experimental platform to test if TAD boundaries restrict lamina-release.

We therefore applied scDam&T-seq to wildtype and ΔCTCF mutant hindlimb cells at E13.5 where lamina-release has normally already occurred (ED Fig. 11b and Table 12). Both wildtype and mutant conditions contained the same cell type proportions as in our larger dataset (ED Fig. 11c and e). Yet, in ΔCTCF mutant limbs, *Kcnj2* was ectopically activated in a *Sox9*-like pattern specifically in chondrocytes (ED Fig. 11d). Thus, scDam&T-seq captures the previously reported *Kcnj2* transcriptional misregulation in mutant ΔCTCF embryos^50,51^.

We further assessed how the TAD boundary deletion impacted genome-lamina interactions. To do so, we quantified the average Dam-LMNB1 signal from LPM-derived cells with distinct *Kcnj2/Sox9* expression levels, namely low-low and low-high (ED Fig. 11e). High-low and high-high cells were excluded as these were not found in sufficient numbers in each condition for analysis. This showed that, in wildtype cells, the *Sox9* TAD is already pre-released and so displays minimal Dam-LMNB1 signal when *Sox9* expression is low and almost no genome-lamina interactions when *Sox9* activates (Fig. 4c). By contrast, *Kcnj2*’s inactive TAD largely retains genome-lamina interactions in all cells, losing only minimal Dam-LMNB1 signal when *Sox9’s* TAD activates.

Nevertheless, these independent LAD dynamics are significantly reduced following TAD fusion. In mutant ΔCTCF limbs, the formerly separate *Kcnj2* TAD region now mirrors the lamina-release of its *Sox9* neighbour (Fig. 4c). Hence, the *Kcnj2/Sox9* CTCF boundary helps enable the independent genome-lamina interactions of their respective TADs.

LADs are thought to arise from the affinity of heterochromatic regions for nuclear envelope proteins^17,18,32,52^. Deleting the ΔCTCF TAD boundary deletion could therefore release the heterochromatic *Kcnj2* region from the lamina in two ways. Ectopic interactions with active *Sox9* enhancers could convert the *Kcnj2* region to a euchromatic state, thereby abolishing its lamina affinity. Alternatively, the *Kcnj2* region could remain heterochromatic in ΔCTCF limbs but nonetheless have its lamina interactions sterically blocked by its intermingling with the euchromatic *Sox9* region.

To test these possibilities, we performed polymer modelling of a simulated *Kcnj2/Sox9* locus where chromatin intermingling and lamina affinity could be varied independently (see Methods and ED Fig. 11f). We fixed the *Kcnj2*’s lamina affinity at its wildtype value and then simulated the CTCF boundary deletion to increase *Kcnj2/Sox9* region intermingling (ED Fig. 11f). This computer model accurately reproduced the separation of the *Kcnj2* and *Sox9* TADs in wildtype limbs and their fusion in ΔCTCF mutants in simulated Hi-C maps (Fig. 4d,e). Crucially, this ectopic intermingling was also sufficient to block the *Kcnj2* region from stably contacting the lamina, despite it having a positive lamina-affinity included in ΔCTCF mutant simulations (Fig. 4f-h and ED Fig. 11g). Time lapse movies of ΔCTCF simulations illustrate why this occurs - greater intermingling with euchromatic *Sox9* chromatin often physically obstructs the *Kcnj2* region from sustaining lamina interactions (Videos 1-2). This suggests that TAD boundaries help set the extent of lamina-release in the limb by preventing intermingling between distinct chromatin states.

## DISCUSSION

We present the first joint *in vivo* single-cell measurements of LADs and transcription and their connection to chromatin accessibility during mouse organogenesis. By moving beyond *in vitro* or population-averaged approaches, we reveal how multipotent limb progenitors dynamically reconfigure chromatin during cell-fate decision making. We find that limb multipotency is characterised by an extensive, synchronous release of genome-lamina interactions at critical developmental loci. Strikingly, this lamina-release often precedes gene activation, even by up to several developmental days. Thus, though lamina-release is likely necessary for transcription, it is not sufficient by itself^23^. Instead, our data support an earlier model^12^ in which early lamina-release licenses loci for subsequent activation, thereby enabling multipotent progenitors to explore diverse potential cell-fate trajectories.

A key insight is that lamina-release frequently encompasses entire TADs, thereby generalising past locus-specific examples^14,30,53^, and coincides with the binding of key limb TFs. This suggests a highly coordinated process in which genes are commissioned together with the surrounding regulatory machinery that coordinates their expression. Our findings therefore contrast previous examples of more localised LAD release at only genes^22^ or isolated regulatory elements^15,29^. Notably, a previous study found that TF binding triggers lamina-release *in vitro*^30^. Together with our observation that putative TF binding coincides with lamina-release, this strongly suggests a causative role for limb TFs in LAD reorganization during limb development. Among these potential TFs are bHLH TFs (TWIST1, TCF4) and homeodomain TFs (e.g. MSX1, PRRX1, PRRX2, LHX2, LHX9) which have established roles in early limb development^34,38,40,45,54^. Intriguingly, these TFs were also recently shown to drive mesoderm differentiation, including in the limb, by cooperatively binding a composite “Coordinator” motif^45^. This raises the possibility that landscape-scale LAD reorganisation requires the concerted action of multiple cooperating TFs.

We also note that many of these limb TFs are themselves encoded by genes located in detaching LAD regions (e.g. *Lhx2, Lhx9, Prrx1, and Twist1*). What’s more, we find these TFs are predicted or confirmed to bind throughout their own encoding gene’s regulatory landscapes. For example, TWIST1 also binds to validated *Twist1* enhancers^55^ distributed throughout the *Twist1* TAD, which itself undergoes lamina-release as accessibility is gained (ED Fig. 10). Together, this suggests a potential autoregulatory feedback between TF function and LAD reorganisation. In turn, this would generate a cascade of LAD-release that unlocks loci for later limb development.

We also demonstrate that TAD boundaries further help restrict the physical extent of lamina- release at the *Sox9*/*Kcnj2* locus by blocking loop extrusion. In doing so, this limits the intermingling between adjacent chromatin regions with distinct lamina-affinities, thereby granting them independent dynamics. This illustrates that the interplay between loop extrusion and genome-lamina interactions can structurally coordinate regulatory landscape function. Nevertheless, this TAD-LAD relationship is not absolute. We frequently observe partial or overlapping lamina-release spreading to neighboring TADs (Fig. 3b and ED Fig. 5c), including at the *Sox9/Kcnj2* locus (Fig. 4c), adding to previous examples of cooperative behaviour between LAD regions^56^. Likewise, our prior work at the *Fat1* locus demonstrates that local promoter/enhancer detachment and activation can occur while most of the surrounding TAD remains lamina-associated^29^. The interplay between chromatin topology and lamina-genome interactions is thus complex and remains challenging to predict.

Our study demonstrates how cells dismantle lamina-heterochromatin interactions and lay the foundations for future cell-fates. A key next step will be to identify the mechanism driving lamina- release. For example, cooperative TF binding may break multivalent genome-lamina interactions by removing the histone modifications that are proposed to build them, such as H3K9me2^32,52,57^. Likewise, this could also be achieved by TFs introducing chromatin decompaction and/or active modifications, such as histone acetylation, that are incompatible with genome-lamina associations^23,25^. It will also be exciting to determine whether this “priming” principle commonly applies across the cell’s collective heterochromatin machinery. This includes H3K9me2, H3K9me3 and polycomb repression, all of which are known to reconfigure during differentiation to govern cell-fates^5,11,58–60^. Finally, we must test whether such stepwise chromatin changes also facilitate cell-fate transitions in other contexts, such as regeneration, oncogenic transformation, and inducible cell reprogramming.

## METHODS

### Materials

#### Data collection

scDam&T-seq data collection was performed in transgenic C2 ESCs^62^ and mice carrying an inducible Dam-LMNB1 transgene, generated in this study (see below). For mice, Dam-LMNB1 animals were bred with CAG-CreER^63^ mice to facilitate induction and, for mutant analyses, also with Sox9-Kcnj2 ΔCTCF^50^ mice. All mouse lines were maintained in a C57Bl.6/J background. For scDam&T-seq ESC samples, a CreERT2 coding sequence was incorporated into the Dam-LMNB1 transgene to facilitate induction in C2 ESCs.

Samples for 10x Multiome data were collected from C57Bl.6/J mice sourced from Inotiv as well as G4 ESCs (XY, 129S6/SvEvTac x C57BL/6Ncr F1 hybrid)^64^.

#### Animal husbandry

All mice were housed in a centrally controlled environment with a 12 h light and 12 h dark cycle, temperature of 20–22.2°C, and humidity of 30–50%. All mice had access to food and water ad libitum. Bedding, food and water were routinely changed. All experiments involving animals were carried out following institutional guidelines as approved by LaGeSo Berlin and following the Directive 2010/63/EU of the European Parliament on the protection of animals used for scientific purposes under licences G0248/13 and G0114/18.

#### Cell culture

Mouse G4 ESCs^64^ (XY, 129S6/SvEvTac x C57BL/6Ncr F1 hybrid) or C2 ESCs^62^ (Cellosurus) were grown as described previously on a mitomycin-inactivated CD1 mouse embryonic fibroblast feeder monolayer on gelatinised dishes at 37°C, 7.5% CO2^64,65^. All ESCs were cultured in ESC medium containing knockout DMEM with 4,5 mg/ml glucose and sodium pyruvate (Gibco, 10829018) supplemented with 15% FCS (PAN-Seratech, 2602P122011), 10 mM Glutamine (Biochrom, K0302), 1x penicillin/streptomycin (Biochrom, A2213B), 1x non-essential amino acids (Gibco, 11140050), 1x nucleosides (Sigma, ES008D), 0.1 mM beta-Mercaptoethanol (Sigma, 805740) and 1000 U/ml LIF (Millipore, ESG1106). Medium was changed every day while ESCs were split every 2-3 days. For cryopreservation, ESCs were frozen at one million cells per cryovial in ESC medium containing 20% FCS and 10% DMSO (Sigma, D2650). ESCs and feeder cells were regularly tested for Mycoplasma contamination (Lonza, MycoAlert detection kit.).

#### Generation of inducible Dam-LMNB1 ESCs and mice

The flippase (FLP)-flippase recognition target (FRT) system was used to introduce Dam-LMNB1 transgene constructs into C2 ESCs^62^. This modified ESC line contains a phosphoglycerate kinase neomycin selection cassette flanked by FRT sites and a promoter- and ATG-less hygromycin cassette targeted downstream of the *Col1A1* locus^62^. 800,000 C2 ESCs were seeded onto a feeder-coated 6-well plate and transfected with 9 μg of targeting construct, 3 μg FLP-encoding vector, 1 μl Lipofectamine LTX Plus reagent (Thermo Fisher Scientific), 20 μl Lipofectamine LTX to a final volume 250 μl with OptiMEM (Thermo Fisher, 11960085). After 24 h, transfected C2 cells were transferred onto hygromycin-resistant DR4 feeders and treated with hygromycin B (final concentration 150 μg/ml) in ES growth medium for 5-10 days. Single clones were then picked and transferred to CD1 feeders in 96-well plates. Plates were subsequently split into triplicates after 2- 3 days, two for freezing and one for DNA harvesting. Following lysis and PCR genotyping to validate integration into *Col1a1* locus, selected clones were expanded from frozen plates after which genotypes were reconfirmed. To establish a mouse line, Dam-LMNB1 C2 ESCs were used to produce animals through diploid aggregation^66^, where genotyping confirmed the presence of the desired mutations. A single founder was bred with C57Bl.6/J wild-type animals to establish the line. A full list of ESC and mouse lines and the plasmids and primers used to generate them can be found in Tables 13-14.

#### Dam-LMNB1 induction

Dam-LMNB1 expression was triggered in transgenic ESC and mouse lines via recombination mediated by a tamoxifen inducible Cre-ERT2. In ESCs, 1 uM of 4-hydroxy tamoxifen (4-OHT) (Sigma, SML1666) was added to the media 7 hours prior to harvesting the cells. In mice, tamoxifen was delivered by intraperitoneal injection. Specifically, an injection solution was prepared containing 45 mg/ml tamoxifen (Sigma-Aldrich, T5648) and 5 mg/ml progesterone (Sigma-Aldrich, P3972) in 5% ethanol and 95% corn oil (Sigma-Aldrich, C8267). The mixture was incubated at 37°C with periodic vortexing until tamoxifen was completely dissolved (∼4-6 h). The solution was then aliquoted into 1.5 ml light-protected Eppendorf tubes and stored at -20 °C. Each tube was thawed as needed and used after 5 min equilibration to room temperature. A 1-ml syringe with a 26-gauge needle was used for intraperitoneal injection. Plug-dated pregnant females were injected intraperitoneally with 200 μl of stock solution (9 mg of tamoxifen and 1 mg of progesterone). Embryos were harvested from injected mothers 17 hours post-injection for data collection and between 8-24 hours post-injections for optimization experiments. Tissues were then prepared as described below.

#### Embryo dissections and processing

Embryos were isolated from the uteri of pregnant mothers between 8.5 and 13.5 days after the plug date. Embryos were dissected on ice in HBBS (Gibco, 14025092) (for 8.5-day embryos) or PBS (Gibco, 14190094) (for 9.5-13.5-day embryos), and staged based on morphology, size, and/or somite number (Tables 1, 8). In all cases, extra-embryonic tissues were removed as thoroughly as possible before further processing. Excess material, such as the chorion or tail, was taken for PCR genotyping, as described previously^50^. Primers used for genotyping can be found in Table 14.

Dissected samples were processed to create single cell suspensions differently depending on stage. Whole E8.5 embryos were pooled by precise somite number and transferred for tissue dissociation in 200 μl TrypLE Express (Gibco, 12604013) at 37 °C. Embryos were then dissociated through trituration by pipetting at 10-20 minutes intervals for a total of 40 minutes. By contrast, tissues from E9.5 to E13.5 embryos were microdissected and pooled if needed. Tissues were then dissociated with Trypsin-EDTA (Thermo Fisher, 25300096) and triturated by pipetting at 37°C at 2-3 minute intervals until no visible clumps remained (∼5-15 minutes). Digestion was immediately halted by addition of 10% FCS/PBS. Following disassociation, all samples were cleared of cell clumps by filtration (Scienceware, 40 μm, Flowmi Cell Strainer) and then pelleted by centrifugation at 300 g for 5 minutes at 4°C.

For scDam&T-seq, cells were resuspended in FACS buffer consisting of 10%FCS/PBS, 10μM ROCK inhibitor (AbMole, M1817), 1μg/ml DAPI (Sigma-Aldrich, D9542) in preparation for single-cell sorting. For 10x multiome processing, cells were resuspended in 1:1 ratio of F1 media (40% FCS/60% DMEM (Gibco, 11960044)) and F2 solution (20%DMSO/40%FCS/40% DMEM) and cryopreserved. For microscopy, embryos were fixed overnight in 4%PFA (Roth, 0335.3)/PBS at 4°C. Cells were then washed the next day in PBS and stored 4°C before histology preparation.

### Genomics

#### scDam&T-seq

##### Single-cell sorting and FACS gating strategy

FACS sorting of live cells was performed on the BD FACS Aria III Cell Sorter System with sample and plate cooling at 4°C. Gating was based on cell size (forward/side scatter), exclusion of doublets, and live-dead staining by DAPI. Induced Dam-LMNB1 cells were further gated by GFP expression, a co-expressed reporter protein, relative to non-induced controls. One cell per well was sorted into hard-shell 384-well plates pre-filled with 5 ul/well mineral oil and 100 nl/well 1.5 uM CELseq2 primers ^31,67^. After sorting, plates were sealed with aluminium seals, centrifuged at 300g for 5 minutes at 4°C, and stored at -80°C until processing.

##### Single-cell Dam&T-seq

Single-cell Dam&T-seq was performed as described^67^ with adaptations. These include reduced reaction volumes and adapter concentrations to improve cost efficiency and achieve optimal ratio between DamID and transcription readouts. Specifically, 50 nl/well of Lysis mix (15 nl 10 mM dNTPs, 15 nl 1:50000 RNA spike-ins, 11.25 nl 1% NP-40, 8.75 nl ultra-pure water) were added to a cumulative volume of 150 nl and plates were incubated at 65°C for 5 minutes. Reverse transcription (RT) of polyadenylated RNA was performed with dispension of 100 nl/well of RT mix (50 nl 5x First Strand buffer, 25 nl 0.1 M DTT, 12.5 nl (0.5U) RNase OUT, 12.5 nl (2.5U) Superscript II) to a cumulative volume of 250 nl, and incubation at 42°C for 1h, 4°C for 5 minutes and 70°C for 10 minutes. Second-strand synthesis (SSS) followed, with dispension of 885 nl/well of SSS mix (227 nl 5x Second Strand buffer, 21.8 nl 10 mM dNTPs, 7.8 nl (0.078U) E. coli DNA Ligase, 29.5 nl (0.295U) DNA polymerase I, 7.4 nl (0.015U) RNase H, 591.5 nl ultra-pure water) to a cumulative volume of 1135 nl, followed by incubation at 16°C for 2 hours. Next, the samples were treated with 250 nl/well of Proteinase K mix (69.3 nl 20 mg/ml Proteinase K, 138.5 nl 10x Cutsmart buffer, 42.3 nl ultra-pure water) to a cumulative volume of 1385 nl, followed by incubation at 50°C for 10h and heat inactivation at 80°C for 20 minutes. Then, 115 nl/well of DpnI mix (15 nl (0.3U) DpnI, 15 nl 10x Cutsmart buffer, 85 nl ultra-pure water) were added to a cumulative volume of 1500 nl, followed by incubation at 37°C for 6 hours and heat inactivation at 80°C for 20 minutes. The content of each well was barcoded with the addition of 50 nl/well of DamID2 adapters to a final concentration of 5 nM, followed directly by dispension of 450 nl/well of Ligation mix (200 nl 10x Ligase buffer, 50 nl (0.25U) T4 Ligase, 200 nl ultra-pure water) to a cumulative final volume of 2000 nl. Finally, plates were incubated at 16°C for 16 hours and heat-inactivated at 65°C for 10 minutes. After each dispension step, the plates were sealed with aluminum plate seals and spun at 2000g for 1-2 minutes at 4°C. The content of each plate was pooled by centrifugation of the 384-well plates on collection plates, and subsequent steps until library preparation were performed as previously described^67^.

##### Library preparation and sequencing

Library preparation for scDam&T-seq was performed as reported previously^67,68^ with minor modifications. Specifically, a maximum of 6 μl of 100-200 ng aRNA was used for RT, followed by 8-10 cycles of library PCR, depending on the amount of input aRNA. Purified libraries were sequenced with Illumina NextSeq 500, Nextseq 2000 or NovaSeq 6000 platform with paired-end 75/75bp or 100/100bp read length.

#### 10x Multiome for single ATAC and gene expression profiling

Cryopreserved single-cell suspensions were used to prepare nuclei according to <u>10x Genomics</u> <u>guidelines</u> with several modifications. Briefly, cells were thawed at 37°C and immediately washed twice with warm ESC media by centrifugation at 400 g for 10 minutes at RT. Pipetting was performed as gently as possible to ensure cell viability. Cells were then transferred onto ice and washed twice with cold 0.04%BSA (Sigma, A2153)/PBS by centrifugation at 400 g for 10 minutes at 4°C. After the final wash, the pellet was thoroughly resuspended in lysis buffer diluent (10 mM Tris-HCl pH 7.4 (Roth, 4855.1; Merck, 100317), 10 mM NaCl (Invitrogen, AM9760G), 3 mM MgCl₂ (Invitrogen, AM9530G), 1% BSA, 1 mM DTT (Roth, 6908.3) in DEPC (Sigma, D5758) water) to generate single cell-suspension. An equal volume of 2x concentrated lysis buffer (20 mM Tris-HCl pH 7.4, 20 mM NaCl, 6 mM MgCl₂, 0.20% Tween-20 (Serva, 37470), 0.20% NP40 substitute (Sigma, 74385), 0.02% Digitonin (Milipore, 30410), 2% BSA, 2 mM DTT, in DEPC water) was added to the cell suspension, mixed by inversion, and left to incubate on ice for 3 minutes (embryonic tissues) or 5 minutes (ESCs). Nuclei were then washed three times with Wash buffer (10 mM Tris-HCl pH 7.4, 10 mM NaCl, 3 mM MgCl₂, 1% BSA, 0.10% Tween-20, 1 mM DTT, in DEPC water) by centrifugation at 500g for 5 minutes at 4°C. Prior to the last centrifugation, samples were cleared of cell clumps by filtration (Scienceware, 40 μM Flowmi Cell Strainer). Nuclei were resuspended in 10x Genomics diluted nucleus buffer (PN-2000207), quantified using a haemocytometer (Neubauer, 0640110) and adjusted to 3,220 nuclei per μl. In total, 5 μl of diluted nuclei were subsequently used for the transposase reaction. All steps were performed on a RNAse-Zap (Sigma, R2020) treated bench. All buffers were supplemented with 1 U/μl RNAse inhibitor (1:1 ratio of Invitrogen, AM2684 and AM2694). The final libraries were sequenced on the Illumina NovaSeq 6000 or NovaSeq X Plus platform with paired-end 50/50bp read length for ATAC modality and 28/90bp or 50/90bp for RNA modality.

### Microscopy

#### Sample preparation

Mouse embryos were fixed overnight in 4%PFA/PBS and subsequently washed twice with PBS. Embryos were then cryoprotected by incubation with 10%, 20% and 30% sucrose (Roth, 9097.1), and embedded into Jung freezing medium (Leica Microsystems) using Disposable Embedding Molds (Polysciences Inc.). Cryoblocks were stored at −80°C. Cryosections with thickness of 16-20 μm were sliced in frontal, sagittal and transverse planes (Leica Microsystems Cryostat), collected on SuperFrost microscopic slides (Roth, SuperFrost Ultra Plus), immediately frozen and stored at −80°C before use.

#### Immunofluorescence

Cryosections were air-dried at RT for 30 min, permeabilized with 0.5% Triton X-100/PBS for 30 minutes and washed with PBS. Primary and secondary Abs were diluted in blocking solution consisting of PBS, 2% BSA, 0.1% saponin, 0.1% Triton X-100. Incubation with Abs was performed in home-made chambers placed in dark wet containers at RT overnight as previously described^69^. For nuclear counterstain, DAPI was added to secondary Ab to a final concentration of 2 µg/ml. Washing after incubation with antibodies was performed with PBS/0.01% Triton X-100 pre-warmed to 37°C, 3 x 30 min each at 37°C. Sections were mounted under coverslips with antifade medium Vectorshield (VectorLabs) and sealed with nailpolish.

The following primary and secondary antibodies were used: (i) rabbit anti-mCherry (1:500, Abcam, ab167453) and donkey anti-rabbit conjugated with DyLight 594 (1:500, JacksonImmuno Research, 711-516-152), (ii) chicken anti-GFP (1:100, Aves Labs, GFP-1020) and donkey anti-chicken conjugated with Alexa 488 (1:500, Dianova, 703-546-155), (iii) mouse anti-V5 (1:100, Invitrogen, R960-25) and donkey anti-mouse conjugated with Alexa 555 (1:500, Invitrogen, A31570).

### Image acquisition

Single optical sections were acquired with TCS SP5 confocal microscope (Leica) using a Plan Apo 63/1.4 NA oil immersion objective and the Leica Application Suite Advanced Fluorescence Software (Leica). For overviews of tissues and organs, Leica Tile Scan Protocol was used that allowed collection of separate fields and their automated stitching.

### Computational Methods

All data analysis was performed with R (v. 4.4.3) unless otherwise indicated.

#### Hi-C data reprocessing

Downloaded Fastq files were processed with the Juicer pipeline^70^ (v1.5.6, CPU version) using bwa^71^ (v0.7.17) for mapping short reads to the reference mm10 genome. Replicates were merged after the mapping, filtering and deduplication steps of the Juicer pipeline. Juicer tools^70^ (v1.7.5) were used to generate binned and KR normalized Hi-C maps from read pairs with MAPQ≥30. TADs were identified by applying TopDom v.0.0.228 on 50-kb binned and KR-normalized maps using a window size of 10^72^. Insulation scores were calculated using Cooltools (https://github.com/open2c/cooltools/tree/v0.4.1).

#### Processing of scDamID and scDam&T-seq data

scDam&T-seq data was largely processed with the pipeline described in Markodimitraki et al. (2020) (www.github.com/KindLab/scDamAndTools). The procedure is described in short below.

### Demultiplexing

All reads are demultiplexed based on the DamID and CEL-Seq2 single-cell barcodes present at the start of R1. Zero mismatches are allowed between the observed barcode and reference. The UMI (Unique Molecular Identifier) sequence, intermingled with the barcode, is appended to the read name.

### DamID data processing

DamID R1 reads are aligned using bowtie2^73^ (v. 2.5.3) with the parameters: ‘--seed 42 --very-sensitive -N 1’ to the mm10 reference genome. The resulting alignments are then converted to UMI-unique GATC counts by matching each alignment to known strand-specific GATC positions in the reference genome. Any reads not matching a known GATC position or have a mapping quality lower than 10 are removed. 4 unique UMIs are allowed per position to account for the maximum number of alleles in a diploid single cell. Finally, counts are binned at 20kb resolution.

### CEL-Seq2 data processing

CEL-Seq2 R2 reads are aligned using hisat2^74^ (v. 2.2.1) with parameters: ‘--mp ‘2,0’ --sp ‘4,0’’ since R1 only contains the sample barcode, UMI and poly-A tail. The mm10 reference genome and the GRCm38 (v. 89) transcript models are used. Alignments are then converted to transcript counts per gene.

### Filtering of DamID data

Samples with <10’000 unique number of GATCs were filtered out (see ED Fig. 1f-g and Table 2). Additionally, genomic bins with less than 1 mappable GATC per kb were removed from the analysis.

#### Single-cell transcription analyses

##### Single-cell transcriptional UMAP

The transcriptional UMAP (Fig. 1b, ED Fig. 2a and c-bottom) was generated from single-cell transcript count tables, from which 13 Gm genes were removed, using Seurat^75^ (v. 5.1.0) . These Gm genes are thought to result from some leakage from the DamID reads and were found as a non-biological source of variability. Cells were filtered based on number of unique genes (>1’000 and <7’500), percentage of mitochondrial transcripts (<10%), percentage of hemoglobin genes (<7%), and the percentage of ERCC spike-in derived reads (<3%).

Genes with counts observed in fewer than five cells were excluded. Data was then normalized using the ‘SCTransform’ command, regressing for ERCC spike-in, mitochondrial, ribosomal and hemoglobin genes as well as for CT010467.1 which was identified as an rRNA contaminant. The UMAP was then generated using the ‘FindVariableFeatures’, ‘RunPCA’ and ‘RunUMAP’ commands using 1:40 PCs. Ribosomal, ERCC, hemoglobin, and mitochondrial genes were additionally removed from the set of variable features.

### Marker gene identification

Markers per cell type were found using the FindMarkers command with parameters ‘min.pct = 0.25’ and ‘logfc.threshold = 0.25’. To generate the heatmap in ED Fig. 2b, the following steps were taken: 1) the top 10 markers per cell type were selected based on average log_2_ fold change (log_2_(FC)), 2) from those genes that appeared in the top 10 list more than 3 cell types were removed, 3) average gene expression was then calculated for the top 10 filtered list of genes, 4) the top 2 genes (based on log_2_(FC)) of this filtered list were selected to label the rows of the heatmap, 5) expression data was then z-score normalized.

### Cell type annotation

Cell types were identified by a supervised clustering approach and known marker genes from available atlases and literature^33,46,76–93^. Briefly, we manually adjusted the resolution parameter towards modest overclustering, and then manually merged adjacent clusters if they highly expressed the same literature-nominated marker genes. We then annotated individual cell-types using at least four literature-nominated marker genes per cell-type label (Table 10). These cell-types were then assigned a broader cell-type group based on common developmental origin, function and/or marker genes (Table 4). Cells in clusters that had only non-biologically significant markers were removed from subsequent analysis.

After cell type annotation, cells that did not pass DamID thresholds were removed from the transcriptional dataset for any downstream analysis.

#### DamID analysis

##### DamID data binarisation

Single-cell count tables were converted to binary contacts, where a value of 1 reflects lamina association and a value of 0 reflects no lamina association. To determine binary contacts, the 20- kb binned counts were first depth normalized (RPKM) and smoothened with a Gaussian kernel (s.d. 40 kb). Next, the observed values were compared with the density of mappable GATCs, which underwent the same smoothing and normalization. Finally, contacts were called when the difference between the observed signal and the control was bigger than 0 for Dam-LMNB1 compared to a mappability control.

##### Contact frequency and *in silico* population profiles

Binary contacts were used to compute Contact frequency (CF) as the fraction of single cells in which a contact with the lamina was observed per genomic bin. *In silico* pseudobulk profiles were also calculated by combining the count data of all single-cell samples per condition (e.g. cell types) and calculating RPKM and log_2_(O/E) similarly to what was described above for single cells.

### LAD calling

LADs depicted in Fig. 1e were calculated from binarised log_2_(O/E) values of each cell type with more than 50 cells, where values >0 are considered to be a LAD. Consecutive LADs with less than 100kb separating them were joined together. Domains classified as LADs in all cell types were defined as cLADs, while those that were never LADs in any cell type were identified as Non-LADs. fLADs represent genomic regions that range from being detached from the lamina in one cell type to being found in a LAD in only a single cell type. Due to the LAD merging step described above, any overlapping LAD types (e.g. cLADs, fLADs, cNon-LADs) were resolved by selecting the larger LAD and discarding the smaller domains.

#### Single-cell DamID analyses

##### Single-cell DamID UMAP

The UMAP based on single-cell LAD data (Fig. 1c,d and ED Fig. 2c-top) was generated similarly as in Cusanovich et al^74^. We first used the single cell binarized log_2_OE Dam-LMNB1 data (smoothed in 20kb bins) and created a bin x cell matrix. From this matrix, we considered only the top 20,000 most variable bins and performed dimensionality reduction using a term frequency-inverse document frequency (‘TF-IDF’) transformation. We then used singular value decomposition (SVD) on the TF-IDF transformed values to generate a lower dimensional representation of the data by only considering the 3rd through 50th dimensions. Dimensions 1 and 2 were excluded because of high correlation with signal-to-noise ratio of the data. Signal-to-noise was calculated as the FRIP score on called LADs of the corresponding tissue. kmeans clustering was used to classify cells into 16 clusters.

##### UMAP and iterative clustering

Each genomic bin that exhibited differences between these clusters with log₂(O/E) > 0 was considered a separate domain, allowing for a greater resolution in the detection of lamina interaction differences between cells. The overlap between binarized log₂(O/E) values of individual cells at these genomic coordinates was then calculated, generating a domain-by-cell matrix. This matrix was subsequently loaded into Signac^94^, and cells with a LAD FRIP score < 0.9 were filtered out. UMAP embedding depicted in Fig. 1c was computed using LSI dimensions 3:35.

For clustering, we applied the Smart Local Moving (SLM) algorithm, which initially identified 8 clusters. To refine the structure further, UMAP and clustering were iteratively applied to each of the larger clusters using the highest resolution that maintained clear separation in the UMAP. This approach resulted in 21 final Dam-LMNB1 clusters.

#### Genomic region analysis

##### Genomic region UMAP and clustering

To further investigate differences in lamina interactions, we extracted the normalized domain x cell matrix from the same Seurat object (see above) and performed dimensionality reduction on genomic regions rather than single cells. This was achieved by first performing PCA and using the first 20 PCs to compute the UMAP depicted in ED Fig. 3b,c. Clustering of genomic regions was computed with the Louvain algorithm using 8 nearest neighbors which resulted in 16 clusters depicted in ED Fig. 3a, c and d.

##### Domain gene expression enrichment

Genes whose TSS overlapped each LAD cluster were extracted. Gene Ontology (GO) enrichment analysis was then performed on each set of genes using the ‘topGO’ package. For each LAD cluster the 6 highest enriched terms were selected. Only GO terms with p-value < 0.001 were considered.

#### Trajectory analysis

##### Cell type selection

Our analyses of mRNA and LAD dynamics in development required the reconstruction of LPM-derived differentiation trajectories in the limb. For this, we carefully evaluated our annotated cell types and their corresponding marker genes in the context of the extensive literature available on limb development (Tables 3-4 and 10). This allowed us to identify 5 trajectories where successive cell types could be confidently identified from E8.5 to E13.5. We note however there are multiple additional unanalysed cell type trajectories that we did not analyse due to insufficient information on their underlying development.

##### Velocity

Velocyto^95^ (v0.17.17) was used for quantification of spliced and unspliced reads from a merged bam file containing the single cell alignments of cells that passed quality thresholds, using the GRCm38 gene model. The resulting loom file was further analysed using scvelo^96^ (v0.3.1). Here, we considered only limb cells derived from the E8.5 lateral plate mesoderm (i.e. of a mesenchymal origin). The PCA and UMAP embeddings used in the analysis were imported from the Seurat object depicted in Fig. 2a. Each cell type had between 69 and 82% spliced reads and between 13 and 26% unspliced reads. Basic preprocessing steps and default steps were followed to estimate velocity. The calculated velocities were then visualized using the ‘scv.pl.velocity_embedding_stream’ function.

### Trajectory inference and differential expression across pseudotime

To construct the limb trajectories, the larger Seurat object was first subsetted into the cell types of each trajectory. PCA and UMAP embeddings were then recalculated and loaded into the R package ‘monocle3’^89^ (v1.3.7) for trajectory inference. Embryonic lateral plate mesoderm was used as the starting point of all trajectories. Differentially expressed gene across pseudotime were calculated using graph-autocorrelation analysis and selecting for genes with q-value< 0.001.

### Differential lamina association across pseudotime

A similar approach to differential gene expression was used to detect differential lamina association across pseudotime, using a gene x cell matrix containing Dam-LMNB1 RPKM values of the 200kb surrounding the gene’s TSS. The RPKM values were then log-normalised using Seurat functions. The resulting Seurat object was further processed with monocle using dimensionality reduction embeddings obtained from the transcriptional output. In this way, a similar pseudotime axis is used to calculate both differential expression and differential Dam-LMNB1 signal. DLI genes over pseudotime were then calculated using graph-autocorrelation analysis and selecting for genes with q-value< 0.001.

### Differential gene expression and differential lamina association across pseudotime

Once genes with differential lamina association and gene expression were identified for each trajectory, they were further filtered by imposing a Moran’s I threshold > 0.07 and a Pearson correlation between LMNB1 and gene expression values lower than 0.1. This ensures analysis of genes whose expression has some level of anti-correlation with their lamina association.

To determine whether a gene was ‘attaching’ or ‘detaching’ the lamina in a trajectory, genes were clustered into 8 clusters based on the kmeans algorithm. To reduce noise, pseudotime was split into 10 bins and averages of expression and lamina association for each cluster were calculated. If the maximum average RNA value occurred after the highest Dam-LMNB1 value, the cluster was classified as ‘detaching’ while if the maximum average RNA value preceded the highest Dam- LMNB1 value, the cluster was classified as ‘attaching’.

The clusters depicted in Fig. 2d-h and ED Fig. 5a were calculated using the k-means method on the Dam-LMNB1 scaled data of the first 6 cell types of the trajectory. Within each cluster, genes were ordered according to the earliest point at which the respective scaled expression data surpasses the 90th percentile.

### Extent of lamina release analysis

For the enrichment shown in ED Fig. 5c, single-cell binarized log_2_(O/E) values were overlapped with TAD domains identified in available E12.5 limb Hi-C data^44^. TADs smaller than 10 20kb genomic bins were not included. After this filtering, all TADs were scaled to the width of the smallest TAD. Per gene, cells were k-means clustered (2 clusters) to result in a ‘at the nuclear lamina’ cluster and a ‘off the nuclear lamina’ cluster. Binarized log_2_(O/E) was then averaged per cluster.

Random forest modelling for ‘uncoupledness score’

To determine the range of the delay between lamina release and gene activation, we focused only on ‘detaching’ genes. To be able to compare results across trajectories, pseudotime was scaled in a 0-1 range. To reduce noise, gene expression data and log_2_(O/E) values per cell were averaged across 15 nearest neighbors over pseudotime using the ‘FindNeighbors’ and the ‘TopNeighbors’ functions from the ‘Seurat’ package^75^. Only genes with at least 5 cells with RNA expression exceeding 0.5 and with LMNB1 log_2_(O/E) above 0 were considered.

In the PI trajectory, two genes representing either a coupled (*Prrx1*) – immediate activation - or uncoupled (*Ptx3*) behaviour – lagged activation – were to train their respective random forest models (ntree=200). Next, for each gene showing differential lamina association and differential expression across pseudotime, gene expression values were predicted using either the coupled or uncoupled model. To evaluate how well real gene expression values matched predicted values computed by each of the models, Root Mean Square Error (RMSE) was calculated. The ‘uncoupledness score’ was defined as the difference between the RMSE using the ‘coupled’ model and the RMSE using the ‘uncoupled’ model. High ‘uncoupledness’ scores indicate that the gene expression of a gene was more accurately predicted using the uncoupled model.

#### Multiome ATAC and Gene expression analysis

##### Transcriptional analysis

Counts for the open chromatin and transcriptional read-outs were obtained by running the cellranger-arc count pipeline (v2.0.0) with default settings. Cells were filtered based on a number of unique genes (>1’000 and <8’000) and percentage of mitochondrial transcripts (<25). However, the total number of cells in the 10x Multiome dataset was very high relative to the scDam&T-seq dataset, particularly for the E8.5 and E9.5 stages. We therefore randomly subsampled the number of 10x Multiome cells to approximately match that of the scDam&T-seq dataset (i.e. allowing for only up to 1000 additional cells per 10x Multiome library).

Genes with counts observed in fewer than 5 cells were excluded and data was normalized using the ‘SCTransform’ command, regressing for mitochondrial genes. Clusters were generated by using ‘FindClusters’ using 1:30 PCs with resolution 9.

### Cell type annotation

Cell types were identified in 10x Multiome datasets as described for the scDam&T-seq dataset (see above). However, 10x Multiome cells that were found in clusters with no scDam&T-seq equivalent were not considered in subsequent analysis. These include visceral endoderm, notochord and allantois, which were presumably excluded in scDam&T-seq due to biases in Dam- LMNB1 transgene expression. A detailed summary of the 10x Multiome annotation is found in Table 11.

### Integration with scDamT dataset

To integrate the 10X multiome and scDam&T-seq datasets, the ‘merge’ command from the ‘Seurat’^75^ package was used, followed by the normalization of the merged dataset using the ‘SCTransform’ command regressing for mitochondrial, ribosomal and hemoglobin genes as well as for the CT010467.1, which was identified as an rRNA contaminant. The UMAP was then generated using the ‘FindVariableFeatures’, ‘RunPCA’ and ‘RunUMAP’ commands using 1:40 PCs. Ribosomal, hemoglobin, and mitochondrial genes were additionally removed from the set of variable features. Following principle component analysis with ‘RunPCA’, the integration was performed by using the ‘IntegrateLayers’ command with anchor-based CCA integration. UMAP of the merged dataset was then calculated using the ‘FindNeighbors’ and ‘RunUMAP’ functions with 1:40 PCs and clusters identified at resolution 2.

To evaluate the integration between the scDam&T-seq and 10x Multiome datasets, the ratio of each separately-annotated cell type was calculated per cluster of the merged dataset.

### Pseudotime calculation and binning

Each specific trajectory was integrated similarly starting from a subsetted dataset that contained the relevant cell types. Trajectory inference was performed using package ‘monocle3’ (v1.3.7) in the same way as for the scDam&T-seq dataset alone.

In order to be able to more readily compare the timings of the Dam-LMNB1 and ATAC read-outs, pseudotime of the integrated dataset was binned in 50 equal intervals. Gene expression, ATAC and Dam-LMNB1 values of the cells within those pseudotime intervals were averaged. The distribution of cell types per interval per technique is depicted in Fig. 3a and ED Fig. 8a. Binned pseudotime is used throughout Fig.3 and ED Fig. 8.

### Gene regulatory network prediction

We used python (v. 3.11.9) package SCENIC+^48^ (v. 1.0a1) to build enhancer-driven gene regulatory networks (GRNs) on the cell types depicted in ED Fig. 8b to include the cell types preceding and the cell types just following the window of lamina release. Briefly, we started by using pycisTopic (v. 2.0a0) to identify candidate enhancers and perform topic modeling. First, a set of consensus peaks was generated using pseudobulk ATAC-seq profiles of each cell type. A count matrix over this set of regions was then generated to be used in topic modelling using default parameters. The model using 80 topics was selected and topics were binarized following recommended methods (‘otsu’, and ‘topn’). In addition, differentially accessible regions (DARs) between cell types were identified using the ‘find_diff_features’ function on imputed data.

Next, pycistarget (v. 1.0a2) was used for motif enrichment analysis on the binarized topics and DARs were calculated by pycisTopic. ‘create_cisTarget_databases’ was used to create custom cistarget databases as recommended. Cistromes derived from the motif enrichment analysis were then merged by TF to generate a final set of TF-region cistromes. Scenic+ was then run using the files generated in previous steps of the pipeline.

### eRegulon and chromatin accessibility across pseudotime

The resulting eRegulons were then used for the downstream analysis. Genomic regions were only considered when the expression of the predicted transcription factor, the expression of the target gene and ATAC signal showed a positive relationship (+/+ eRegulons).

Only cells with more than 1000 unique genomic regions detected were considered. ATAC data was normalized with the Signac package^94^ (v. 1.14.0) using latent semantic indexing (LSI) using functions ‘FindTopFeatures’, ‘RunTFIDF’ and ‘RunSVD’. The resulting normalized accessibility data was used in downstream analyses.

Single-cell LMNB1 log_2_(O/E) and RPKM values from the scDam&T-seq dataset were averaged in the 200kb window surrounding each genomic region. Differential lamina association of genomic regions was performed with ‘monocle3’ in the same way as for the gene-centric analysis, using Dam-LMNB1 RPKM enrichment at the genomic regions with the dimensionality reduction embeddings from the transcriptional output.

#### ΔCTCF mutant scDam&T-seq analysis

##### Transcription-based UMAP

The transcriptional UMAP (ED Fig. 11c) were generated similarly to the large dataset depicted in Fig. 1b, using Seurat^75^ (v. 5.1.0). Cells were filtered based on number of unique genes (>1’000 and <6’000), percentage of mitochondrial transcripts (<10%), percentage of hemoglobin genes (<7%) and the percentage of ERCC spike-in derived reads (<3%). Normalization and dimensionality reduction was performed similarly as for the large dataset but using 1:30 PCs. In addition, having identified a cluster enriched with cell cycle genes, we additionally regressed out cell cycle effects using ‘SCTransform’ on cell cycle scores obtained from the ‘CellCycleScoring’ function from Seurat.

Single-cell gene expression levels of *Kcnj2* and *Sox9* were defined as ‘high’ if the value in the ‘scale.data’ slot of the Seurat object was higher than 1 and ‘low’ if lower than 1.

### Cell type annotation (mutant dataset)

Cell types were identified for the mutant scDam&T-seq dataset as described previously for our wildtype analysis (see above). We then assigned cells to four major cell lineages at this stage of limb development based on common developmental origin, function and/or marker genes. To validate our approach these lineages were then projected onto scDam&T-seq dataset mRNA UMAP using Seurat label transfer (ED Fig. 11c).

### Polymer modelling

Polymer modelling was performed using a combination of previously published methods. Specifically, we used the Strings and Binders Switch (SBS) polymer model^97^ combined with Loop-Extrusion^98,99^ (LE), in which interaction of binders with a polymer and extrusion of extruding factors along the polymer act simultaneously^100^. In addition, the polymer can interact with a simulated nuclear lamina^29^. We applied this combined model to simulate the 3D structure of *Sox9* and *Kcnj2* TADs with flanking regions (chr11:109860000-113710000; mm10).

#### String polymer

The SBS model simulates a chromatin filament as a string with *N* beads, equipped with binding sites for specific interacting molecules (binders). Binder concentration (*c*) and bead-binder interaction affinity (*E*_*i*nt_) are the system control parameters, as a coil-globule transition occurs when they are above a threshold^97,101^.

In this study, a uniform homo-polymer with *N* = 770 beads has been considered, with affinity and concentration taken above transition threshold and kept constant for the sake of simplicity. Genomic content of each bead is therefore 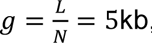, where size of the simulated genomic region is *L* = 3.85 Mb.

Published CTCF ChIP-seq data from E12.5 limbs^61^ was used to define CTCF anchor points following an established approach^100,102^. Specifically, ChIP-seq data binned at 5kb resolution - matching the size of a single polymer bead - was used to call CTCF peaks. These CTCF peaks were then defined as CTCF anchor points with binding probability proportional to signal strength. CTCF orientation was defined using a standard motif search for the CTCF binding motif (from JASPAR database), using the FIMO tool from the MEME suite (https://meme-suite.org/meme/). From this polymer model, we obtain a set of 3D structures representing chromatin conformations in E13.5 limb through standard Molecular Dynamics (MD) simulations^100^.

#### Molecular Dynamics simulations

To generate an ensemble of 3D structures representing the *Sox9-Kcnj2* locus in E13.5 limb, we perform extensive Molecular Dynamics (MD) simulations^100^. For simplicity, bead and binders have the same diameter (*σ* = 1) and mass (*m* = 1) in dimensionless units. To take into account the hard-core repulsion between the particles, we use a purely repulsive truncated Lennard-Jones (LJ) potential. Interaction between beads and binders is modelled with an attractive, short-range LJ potential [inlinewith distance cutoff *R*_*int*_ = 1.3*σ* and regulating the interaction affinity, given by the minimum of the LJ potential and taken as *EE*_*int*_ = 3𝐾𝐾_𝐵𝐵_𝑇𝑇. Consecutive beads of the polymer are linked by FENE bonds^103^ with standard parameters (length *R*_0_ = 1.6*σ* and spring constant 𝐾𝐾_𝐹𝐹𝐹𝐹*N*𝐹𝐹_ = 30𝐾𝐾_𝐵𝐵_𝑇𝑇/*σ*^2^). Positions of beads and binders follow Brownian dynamics and evolve according to the standard Langevin equation^104^ with temperature 𝑇𝑇 = 1, a friction coefficient 𝜁𝜁 = 0.5 and an integration time step 𝛥𝛥𝛥𝛥 = 0.01 (dimensionless units). The initial configuration of the polymer is set as a Self-Avoiding-Walk and the binders are randomly located in the simulation box (linear size 𝐷𝐷 = 50*σ*), then the system is equilibrated up to approximately 10^8^ timesteps. From each model, we perform at least 25 independent simulations in which polymer configurations are sampled every 5*10^4^ timestep once equilibrium is reached.

#### Loop extrusion

Loop-extrusion process is simulated largely as previously described^105^ and is integrated in the above MD simulations where phase-separation and loop extrusion act simultaneously. Specifically, loop extruding factors are modelled as harmonic springs with elastic constant 𝐾_𝑠𝑠𝑠𝑠𝑟𝑟*iiii*𝑠𝑠_ =10𝐾𝐾_𝐵𝐵_𝑇𝑇/*σ*^2^and equilibrium distance 𝑟𝑟_𝑒𝑒𝑒𝑒_ =1.1. Extruding factors slide along the polymer every 500 MD timesteps by moving the spring from the bead pair (*i*, 𝑗) to (*ii* −1, 𝑗𝑗 +1), where *ii*<𝑗𝑗. Extruders can stochastically detach from the polymer with a rate 𝑘𝑘_𝑜𝑜𝑜𝑜𝑜𝑜_, which is related to the processivity through the relation 𝑝𝑝𝑟𝑟𝑝𝑝*c* = 2𝑔𝑔/𝑘𝑘_𝑜𝑜𝑜𝑜𝑜𝑜_. 𝑔𝑔 being the genomic content per bead defined previously^102^.

In our simulations, we considered processivity values in the range of 1.0-1.5Mb to ensure formation of long-range loops and average extruder separation in the range of 250-600 kb^99,105^. When an extruder detaches, a new one is replaced along the polymer in a random position, so as to keep a constant number of extruders. An extruder halts its motion when it meets an oppositely directed CTCF anchor or when it meets another extruder during the sliding, since they cannot pass through each other.

#### Nuclear envelope interactions

To model the nuclear lamina, we use a spherical wall of radius *R* within the simulation box. Polymer beads can attractively interact with lamina though a short range, truncated LJ potential with affinity

*EE*_*N*𝐹𝐹_ ranging from 0.0𝐾𝐾_𝐵𝐵_𝑇𝑇 to 1.4𝐾𝐾_𝐵𝐵_𝑇𝑇 and cutoff distance 𝑟𝑟_*ccii*𝑜𝑜𝑜𝑜𝑜𝑜_ = 2.5*σ*. This is around the lamina- polymer adsorption affinity (around 1.2𝐾𝐾_𝐵𝐵_𝑇𝑇) previously found for similar polymer models^29^. Alternatively, beads interact with the lamina only through a purely repulsive LJ potential which models a mere steric interaction. The lamina sphere radius is set to *R* = 44*σ*.

To determine whether each polymer bead interacts attractively or repulsively with the lamina, we used Dam-LMNB1 signal data collected in this study. Beads with signal values above a threshold— set at the upper tertile of the signal distribution within the modelled locus—were designated as attractively interacting with the lamina.

In our simulations, we first simulate the SBS+LE (polymer+binders+LE) system for 1.5*10^7^ timesteps, sufficient to let the system reach equilibrium^101,106^. Then, we introduce the lamina and in order to ensure the complete interaction of the polymer with the lamina. The system is then equilibrated up to another 8.5*10^7^ timesteps. Simulations have been performed with the HOOMD-Blue package^107^.

#### Calculation of contact maps, lamina association and polymer intermingling

Contact maps are computed as previously described^101,106^. For each polymer configuration, we first measure the Euclidean distance 𝑟𝑟_*iiii*_ between any two beads *ii* and 𝑗𝑗. If the distance is lower than threshold 𝐴𝐴*σ*, the beads are considered in contact. In this study 𝐴𝐴 is taken as 3.5. We have also verified that similar numbers return analogous results. To obtain a frequency map, we then aggregate all the considered configurations sampled at equilibrium from independent simulations. A dimensionless mapping factor is then used to map model maps to experimental numeric range.

The polymer’s association with the lamina (shown in Fig. 4) is computed as the frequency of contact between the polymer and the modelled lamina wall. For this, we first compute Euclidean distance from the lamina for each polymer bead, given by the relation 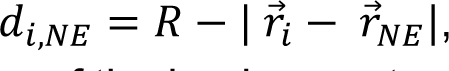, where *R* is the lamina radius, 𝑟𝑟⃗_*ii*_ is the position of bead *ii* and 𝑟𝑟⃗_*N*𝐹𝐹_ is the position of the lamina center. The bead is considered in contact if 𝑑𝑑_*ii*,*N*𝐹𝐹_ < 𝐵𝐵. In this study 𝐵𝐵 is taken as 5. We have also verified that similar numbers return analogous results. We then aggregate the profiles of each polymer configuration to obtain an association frequency. Standard deviations of the profiles are estimated using 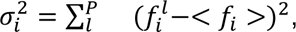, where 𝑃𝑃 is the total number of independent simulations, 𝑓𝑓^𝑙𝑙^ is the association frequency of bead *ii* computed over the sampled configurations within 𝑙𝑙 simulation and < 𝑓𝑓_*ii*_ > is the average frequency over the 𝑃𝑃 simulations.

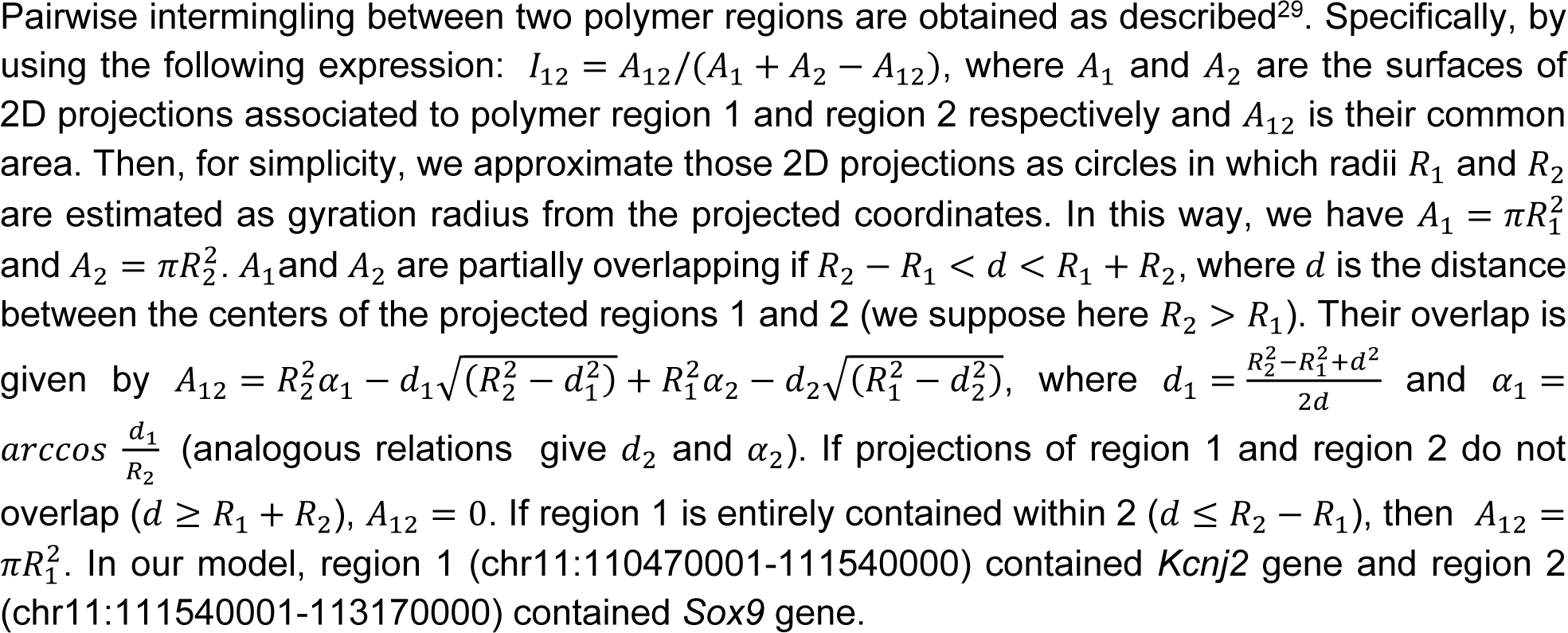

2D contour plots have been generated using the function *kdeplot* from the Python *seaborn* package on the plane showing the intermingling between *Sox9* and *Kcnj2* regions (x-axis) and the fraction of *Kcnj2* region associated to the NE (y-axis), with data points represented by each single polymer configuration.

#### Models of mutants

Simulations of polymer models with deletions of CTCF binding sites (ΔCTCF) are conducted in the following manner. Essentially, the mutation is replicated *in-silico* on the polymer model representing the wildtype condition by setting the probability of CTCF anchor points, which correspond to the deleted CTCF sites to zero^50,61^. The parameters for loop extrusion, i.e. the density of extruders and processivity, along with the affinities between polymer beads and the lamina, are maintained as they are in the wildtype condition. Three-dimensional trajectories are then created through independent molecular MD simulations, as outlined previously.

#### Polymer graphics

The 3D snapshots of the polymer depicted in Fig. 4 are taken from actual MD simulations, in which the *Kcnj2* and *Sox9* regions are distinctly colored. A segment of the simulated nuclear envelope is illustrated as a thick spherical wall, colored to match a FISH image^29^. To better illustrate the spatial relationship between the polymer and the lamina, each image is displayed from a consistent viewpoint utilizing a geometrically calibrated 3D rotation matrix. For enhanced visual clarity, the polymers are represented in a simplified form using a smooth third-order polynomial spline that passes through the coordinates of the polymer beads.

## DATA AVAILABILITY

All data reported in this paper will be shared upon request. Sequencing data generated in this study are available at the NCBI Gene Expression Omnibus (GEO): GSE293955 and GSE293961. Selected processed tracks are also available for visual inspection in UCSC (https://genome-euro.ucsc.edu/s/Chudzik/Chudzik_et_al_2025b). Published TWIST1^45^ and CTCF^61^ ChIPseq was obtained from GSE230316 and GSE84793 respectively. Limb Hi-C^44^ and *Kcnj2*/*Sox9* capture Hi-C^50^ were obtained from GSE275848 and GSE125294, respectively.

## CODE AVAILABILITY

All custom code generated as part of this study is available on request.

## ACKNOWLEDGEMENTS

We thank Corinna Schwichtenberg and Katija Zill (MPIMG Animal Facility) for mouse line maintenance and tamoxifen injections; Ute Fischer and Asita Stiege for genotyping assistance; Alexandra L. Mattei and Maria Walther for training in early mouse embryo dissociation; Claudia Giesecke-Thiel (MPIMG FACs facility) for FACs support with input from Reinier van der Linden (Hubrecht FACS facility); the MPIMG Transgenic unit for aggregations; the MPIMG and MDC, genomics facilities and the Utrecht Sequencing Facility for NGS sequencing; and Robin van der Weide and Pim Rullens for valuable input regarding computational analyses. We also thank Daniel Ibrahim, Guilaume Andrey as well as the Robson and Kind labs for critically reading the manuscript. K.C. was supported by the Deutsche Forschungsgemeinschaft (DFG) International Research Training Group (IRTG2403). I.G. was supported by the Swiss National Science Fund (grant no. P400PB_186758) and an NWO-ENW Veni grant (VI.Veni.202.073). The Utrecht Sequencing Facility is subsidized by the University Medical Center Utrecht and the Netherlands X-omics Initiative (NWO project no. 184.034.019). The lab of M.I.R. was supported by EMBO (ALTF1554-2016), Wellcome Trust (206475/Z/17/Z) and DFG grants (project # 556274722 and FOR 2841, project # 400728090) to M.I.R.. The lab of J.K. was supported by an ERC Consolidator grant (ERC-CoG 1010002885-FateID), a Nederlandse Organisatie voor Wetenschappelijk Onderzoek (NWO) VIDI grant (016.161.339) and and VICI grant (232.033) to J.K.. The lab of S.M. was supported by a grant from the Deutsche Forschungsgemeinschaft (DFG) (SPP 2202/1) and by an ERC Advanced grant (ERC-ADG 101054341 GenRevo) to S.M..

## CONTRIBUTIONS

M.I.R. and J.K. conceived the study with input from S.M.. M.I.R. and K.C. produced Dam-LMNB1 constructs and transgenic ESC/mouse lines. I.S. performed immunofluorescence staining. K.C., M.I.R., S.K., I.G. performed preliminary validations of transgenic lines. K.C. performed all sample collections and cell sorting with support from A.R. and M.I.R.. S.K. optimized and performed all scDam&T-seq experiments and library preparation, with support from I.G.. K.C. performed 10x Multiome with assistance from M.S.. I.G. performed all scDam&T-seq and 10x Multiome computational analyses. I.G., K.C., S.K. and M.I.R. curated the resulting datasets. A.R. reprocessed Hi-C data. A.A. and A.M.C. developed and performed polymer modeling with input from M.N.. K.C., M.I.R., and I.G. prepared figures. M.I.R. wrote the manuscript with input from all authors.

## CORRESPONDING AUTHOR

Correspondence to Stefan Mundlos, Jop Kind, and Michael I. Robson.

## ETHICS DECLARATIONS

The authors declare no competing interests.

## EXTENDED DATA FIGURES

**Extended Data Figure 1.**
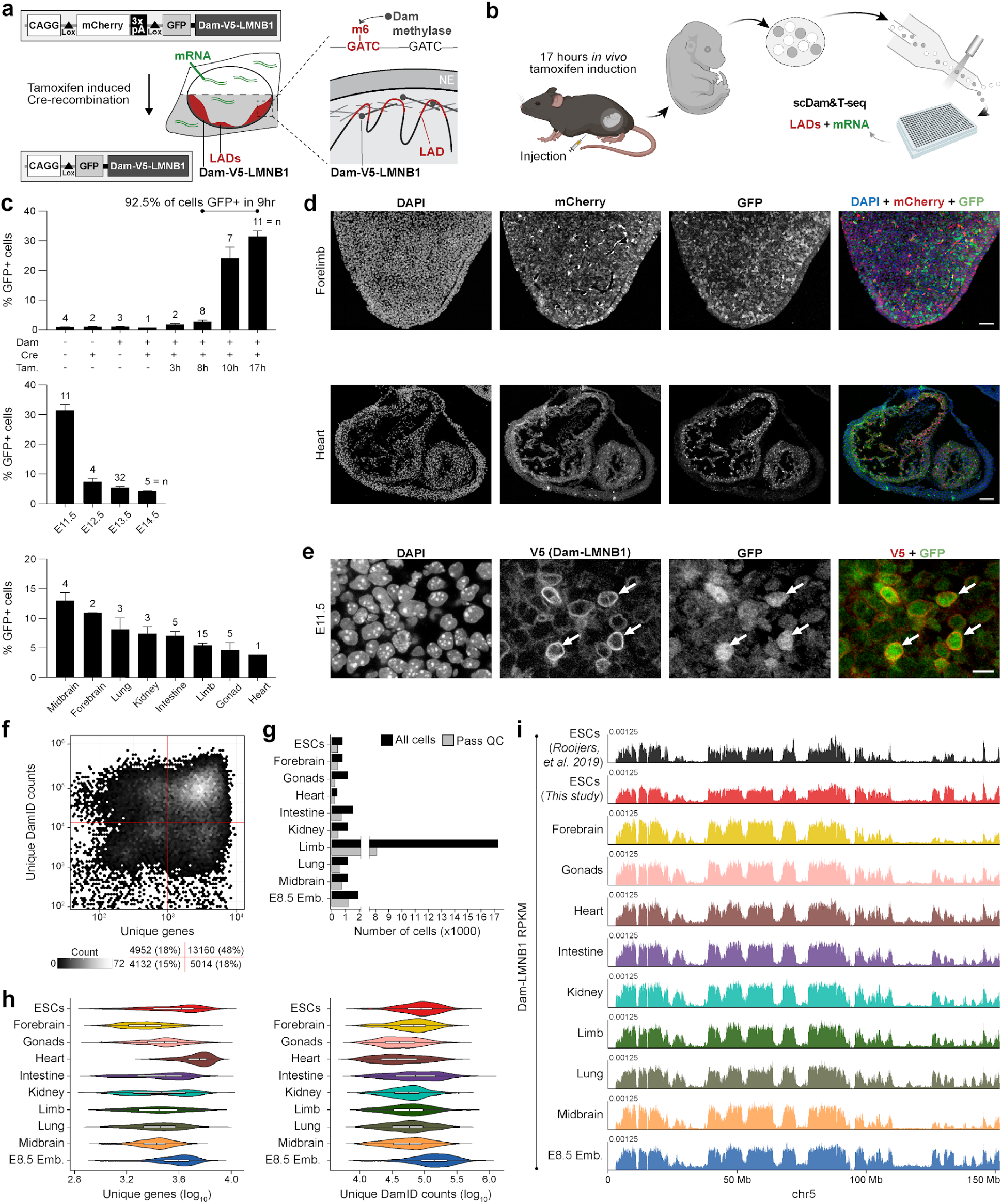
***In vivo* scDam&T-seq jointly profiles single cell LADs and mRNA during mouse embryogenesis. a-b**, Schematic of Dam-LMNB1 mouse allele design (a) and sample collection workflow (b). **c**, FACS analysis of Dam-LMNB1 allele induction over time (top), developmental stages (middle), or between tissues (bottom). 92.5% of cells express GFP from 8 hours after tamoxifen injection. By sampling tissues 17 hours after tamoxifen injection, this therefore gives a ∼9 hour time resolution in which LADs are labelled *in vivo*. **d-e**, Immunofluorescence of tissue sections of induced E11.5 embryos. Scale bars: 100 µm (d - top), 50 µm (d - bottom), 10 µm (e). **f-g**, Quantification of cells that passed quality-control filtering either in all cells (f) or per tissue (g). **h**, Quantification of unique CEL-seq genes and DamID counts detected per cell per tissue, after thresholds in (f) have been applied (ESCs, n = 314; Forebrain, n = 364; Gonads, n = 218; Heart, n = 107; Intestine, n = 668; Kidney, n = 446; Limb, n = 6864; Lung, n = 583; Midbrain, n = 614; E8.5 Emb., n = 1152). **i**, Average Dam-LMNB1 RPKM tracks per indicated tissue. Data from reference ESC track is reprocessed from Rooijers, et al. 2019. See also Figure 1.

**Extended Data Figure 2.**
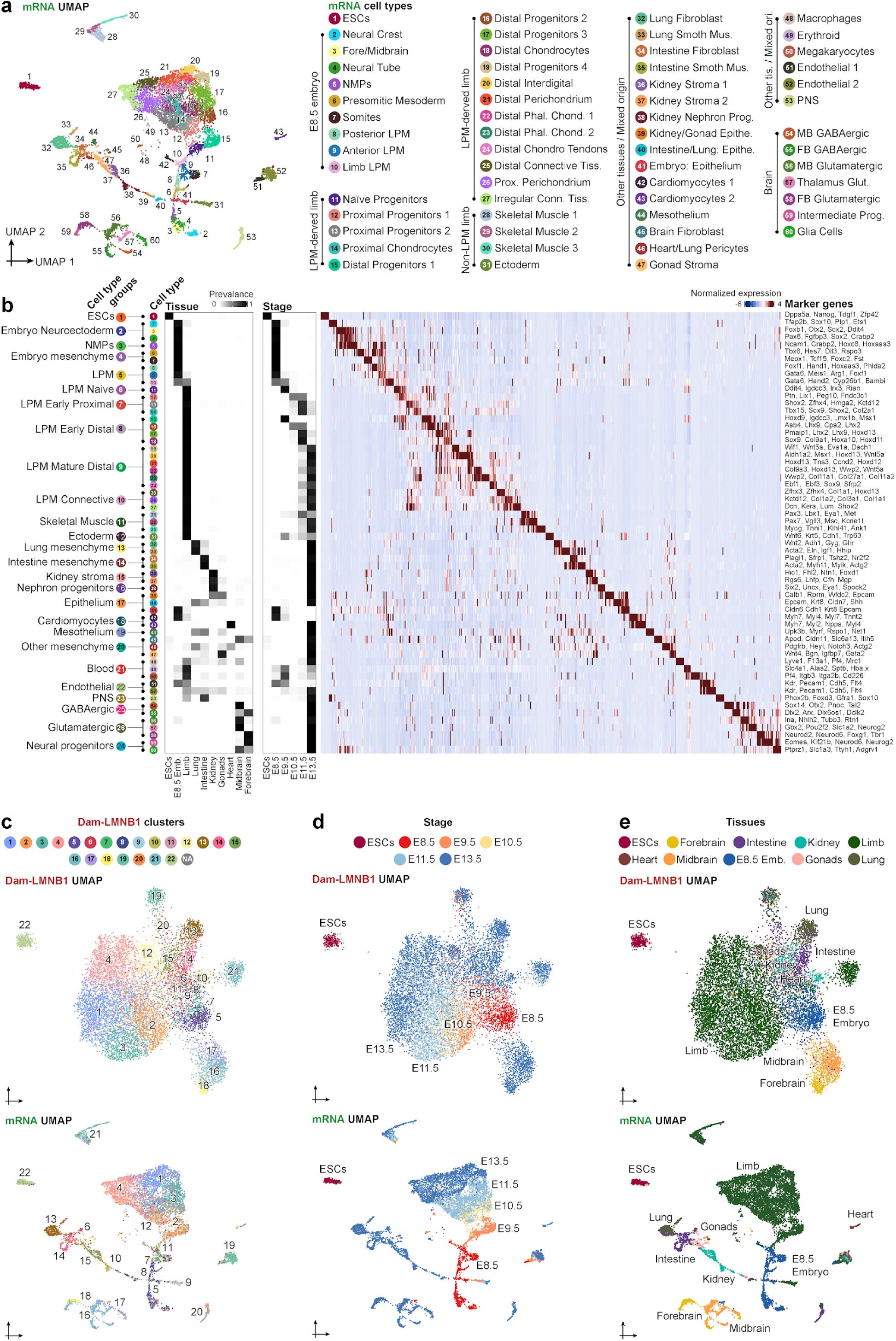
Summary of cell type annotation and additional comparisons of mRNA and LAD clustering. a, UMAP visualization of single cells with coordinates defined by mRNA. Colors and numbers correspond to 60 annotated cell types. **b**, Heatmap with example marker gene expression across mRNA cell types. Relationships of mRNA cell types to broader cell type groups, tissues, and developmental stages are also shown. Prevalence indicates the frequency of a given cell type across the respective tissues and developmental stages. Additional marker genes and annotation information can be found in Tables 3-4. **c-e**, UMAP visualizations of single cells from sampled tissues with coordinates defined by Dam-LMNB1 signal (top) or mRNA (bottom). Colors and labels correspond to Dam-LMNB1 cell clusters (c), embryonic stages (d), or tissues (e). See also Figure 1 and Tables 3-4.

**Extended Data Figure 3.**
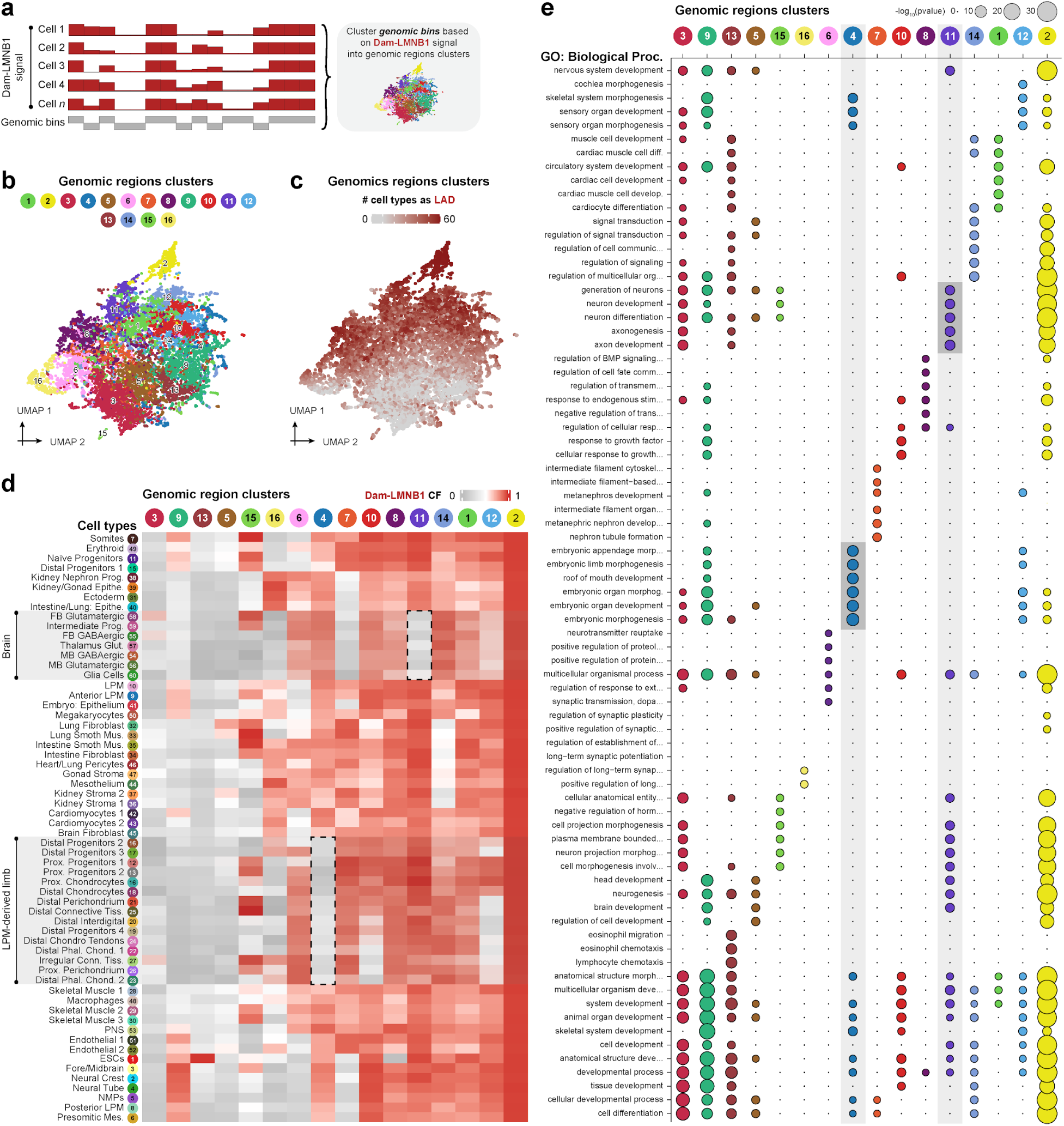
Facultative LADs have cell type specific behaviours and contain genes with relevant developmental functions. a, Strategy to cluster genomic bins by their variation in Dam-LMNB1 signal across single cells (n = 11389). **b**, UMAP visualization of genomic regions with coordinates defined by their Dam-LMNB1 signal across 60 annotated cell types. Colors and numbers correspond to 16 identified clusters of LAD genomic regions. **c**, Matching UMAP colored by the number of cell types in which a given genomic region interacts with the lamina. **d**, Heatmap of average lamina contact frequency (CF) of LAD genomic region clusters across 60 annotated cell types. CF represents the fraction of cells for which a genomic bin contacts the lamina. Dashed boxes highlight example region classes with behaviors in specific cell types. **e**, Enrichment of GO (biological process) filtered for development-related terms for each LAD- region cluster. Vertical grey boxes highlight two example LAD genomic region classes with enrichment of relevant cell type specific GO-terms. See also Figure 1.

**Extended Data Figure 4.**
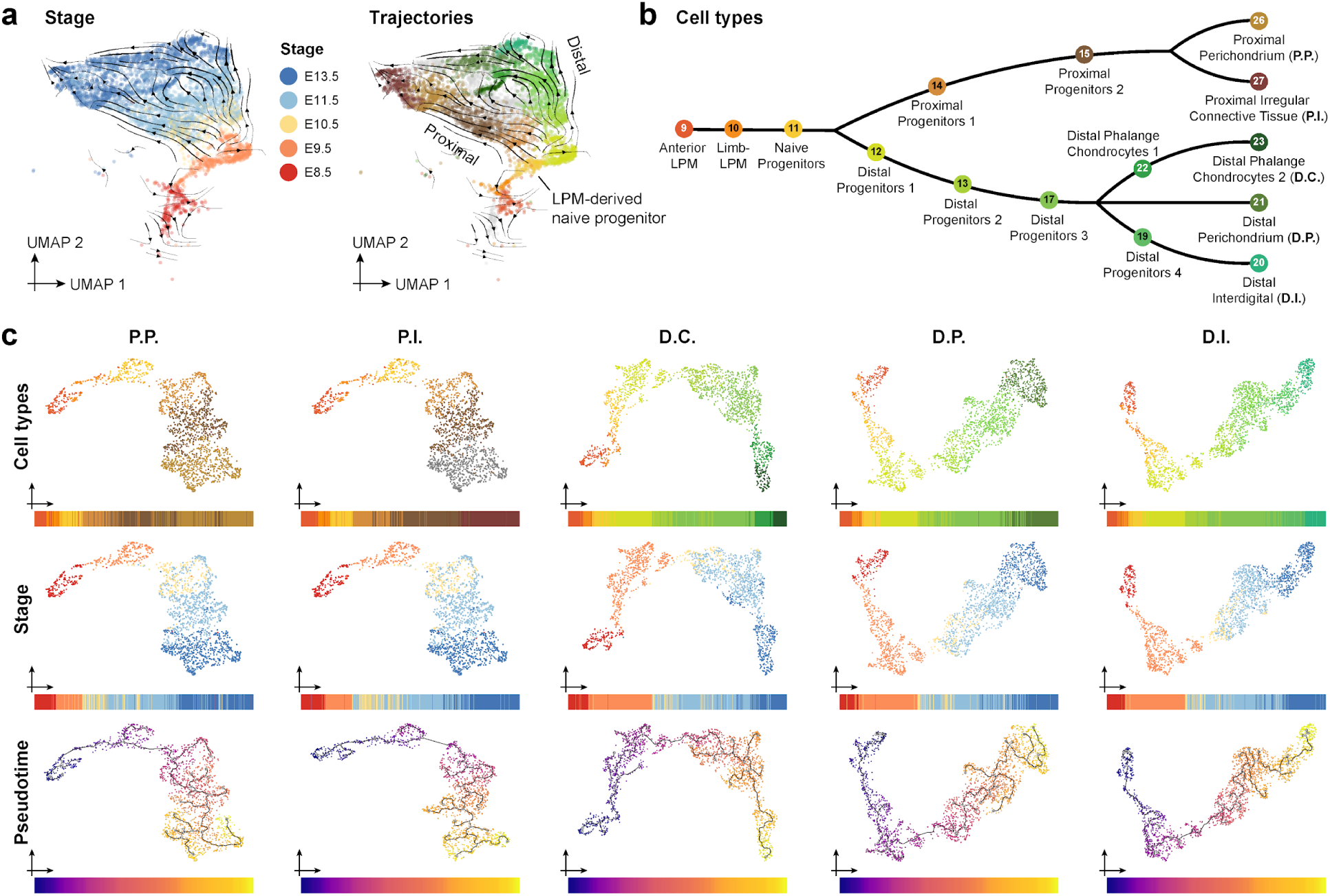
Reconstruction of limb developmental trajectories in pseudotime. **a**, UMAP of LPM-derived limb cell types, with arrows representing RNA velocities calculated using the transcription readout of scDam&T-seq (see Methods). Cells are coloured by embryonic stage (left) or cell types of selected trajectories (right). Arrows indicate the inferred direction of cell differentiation. **b**, Schematic representation of cell type distribution across a subset of five limb developmental trajectories that were selected for analysis. **c**, UMAP of limb developmental trajectories inferred with Monocle (P.P., n = 1772; P.I., n = 1598; D.C., n = 1966; D.P., n = 2116; D.I., n = 2109). Colors correspond to cell type (top), stage (middle), or pseudotime score (bottom) of each cell. See also Figure 2.

**Extended Data Figure 5.**
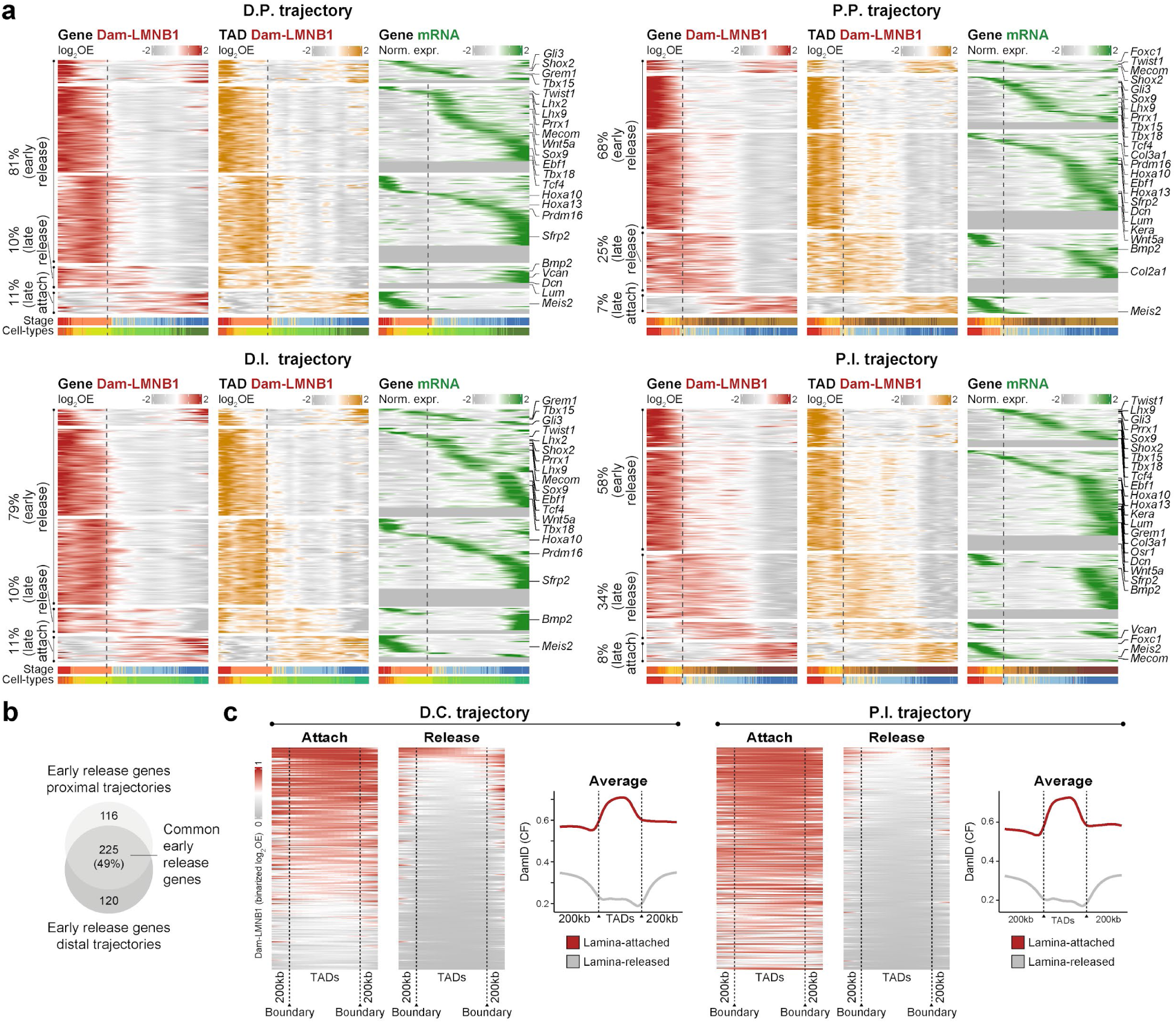
Dam-LMNB1 and mRNA dynamics over pseudotime trajectories for developmental limb genes and TADs. **a**, Heatmap of changing Dam-LMNB1 or mRNA signal for LAD-linked genes along pseudotime in indicated trajectories. Dam-LMNB1 signal is plotted for the 200kb bin containing each gene (red) or their surrounding TAD (orange). Genes are included if they exhibit DLI and DE in at least one trajectory. Example known limb genes are also highlighted. **b,** Fraction of genes that display early lamina release in proximal trajectories and/or distal trajectories. **c,** Heatmaps showing average binarized log_2_OE Dam-LMNB1 values across TADs of individual LAD-linked genes when they are lamina-attached or -released. Signal is plotted over size-normalized TADs and 200 kb of flanking chromatin. Average Dam-LMNB1 quantification for all genes is also shown (right panel). See also Figure 2.

**Extended Data Figure 6.**
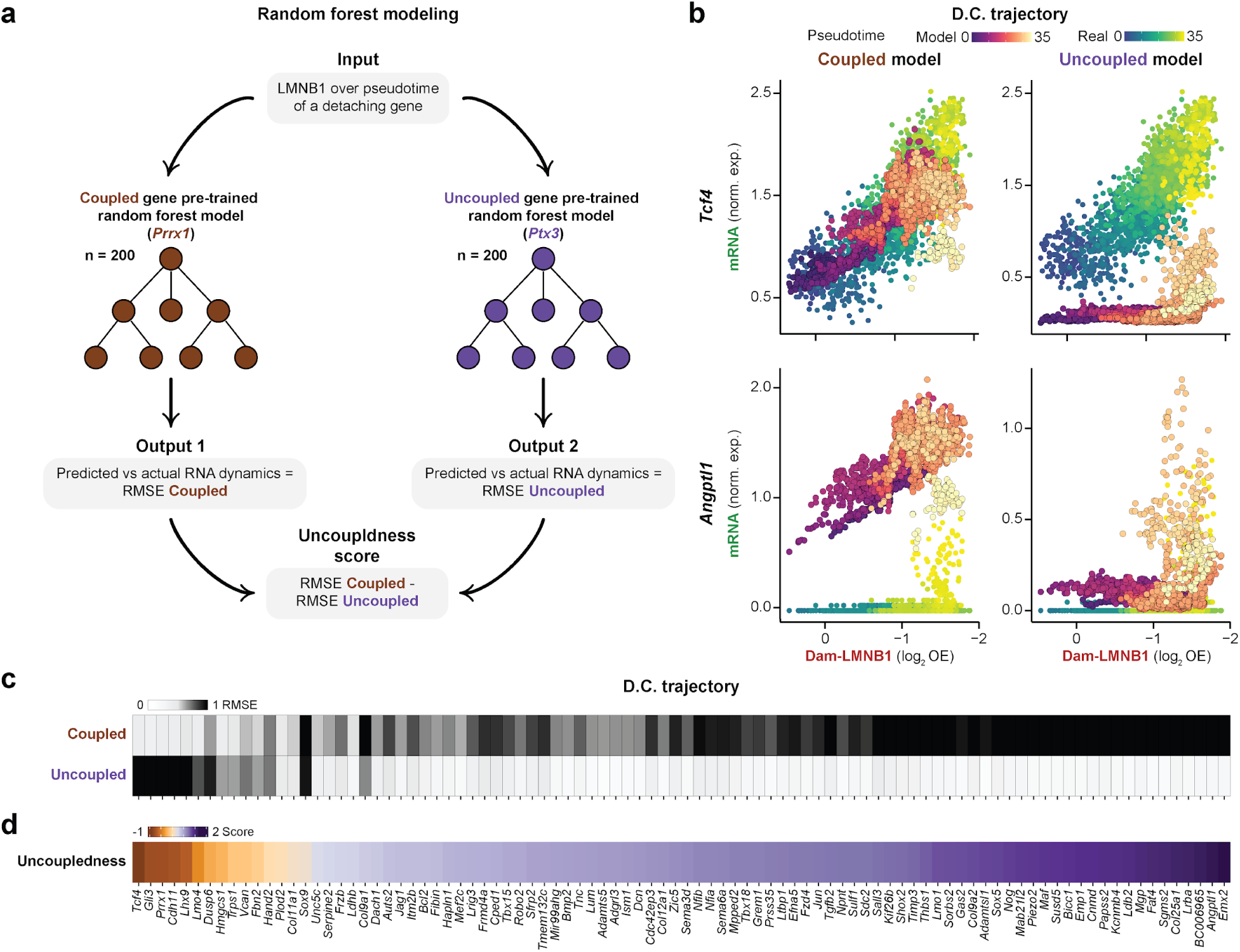
Random forest modeling of Dam-LMNB1 and mRNA dynamics over pseudotime. **a**, Schematic of random forest approach to calculate “uncoupledness score”. Representative genes were used to train random forest models to recognise coupled (*Prrx1*) or uncoupled (*Ptx3*) changes in Dam-LMNB1 and mRNA over pseudotime. **b**, Scatter plots of single- cell Dam-LMNB1 and mRNA signals for two example genes in the D.C. trajectory. Each cell is represented twice in the scatterplots, once with values colored according to real pseudotime, and once with predicted pseudotime values from the coupled or uncoupled model. **c**, Root mean square error (RMSE) using either the values predicted by the coupled (left) or uncoupled (right) model for all detaching LAD-linked genes in D.C. trajectory. **d**, Uncoupledness scores for all detaching LAD-linked genes in the D.C. trajectory. The uncoupledness score is calculated by each gene by subtracting the RMSE for the coupled model predictions from the RMSE from the uncoupled model predictions. Data for the four remaining trajectories are provided in Table 6. See also Figure 2.

**Extended Data Figure 7.**
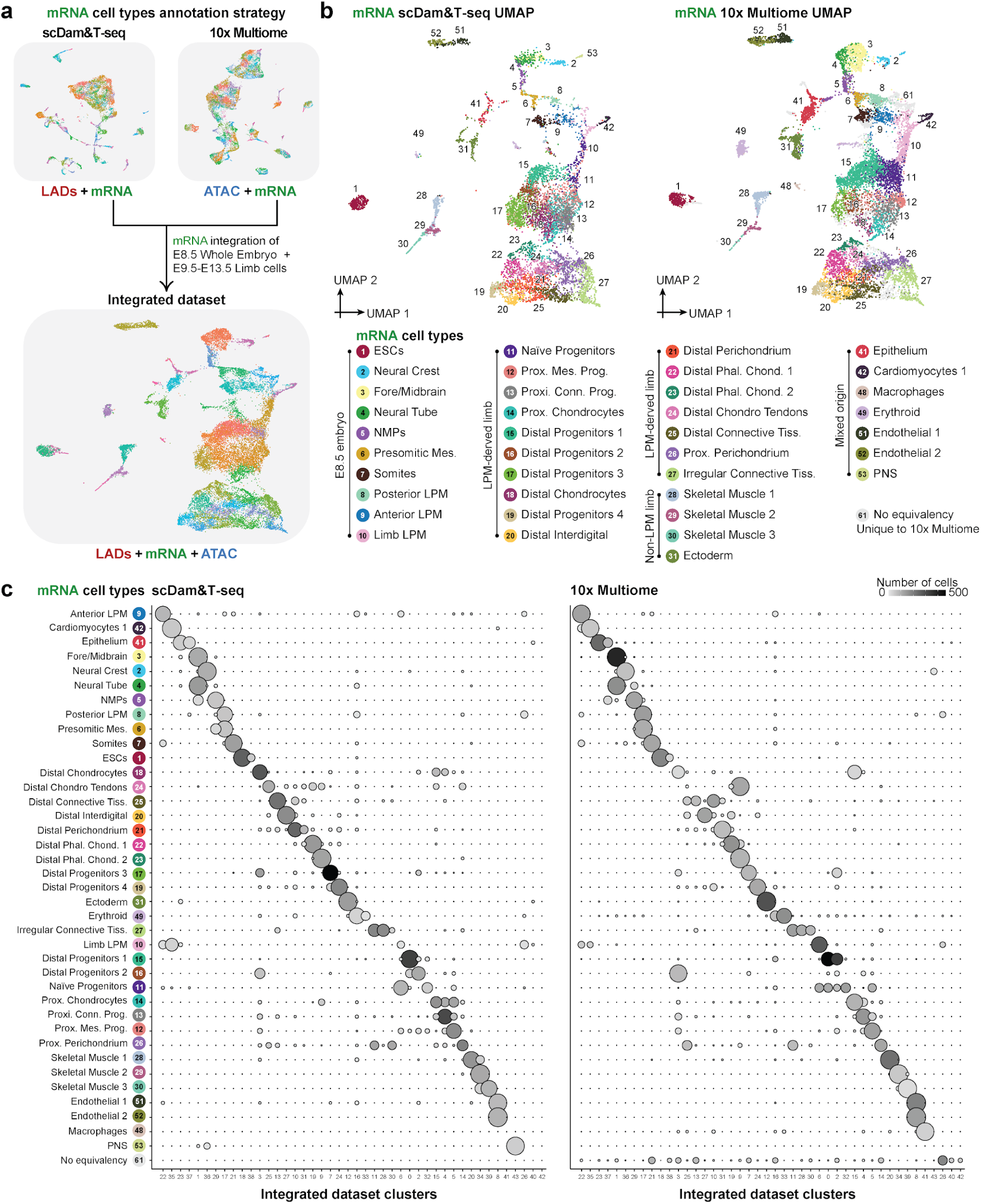
Summary of cell type annotation in the integrated 10x Multiome and scDam&T-seq dataset. **a**, Strategy to integrate cells from scDam&T-seq and 10x Multiome datasets using the common mRNA readout of both technologies. **b**, Integrated UMAP of single cells derived from both scDam&T-seq and 10x Multiome with coordinates defined by the common mRNA readout of both assays. Additional marker genes and annotation information can be found in Tables 10-11. **c**, Dotplot showing the frequency of cells from each assay that are found in clusters identified in the integrated dataset. See also Figure 3.

**Extended Data Figure 8.**
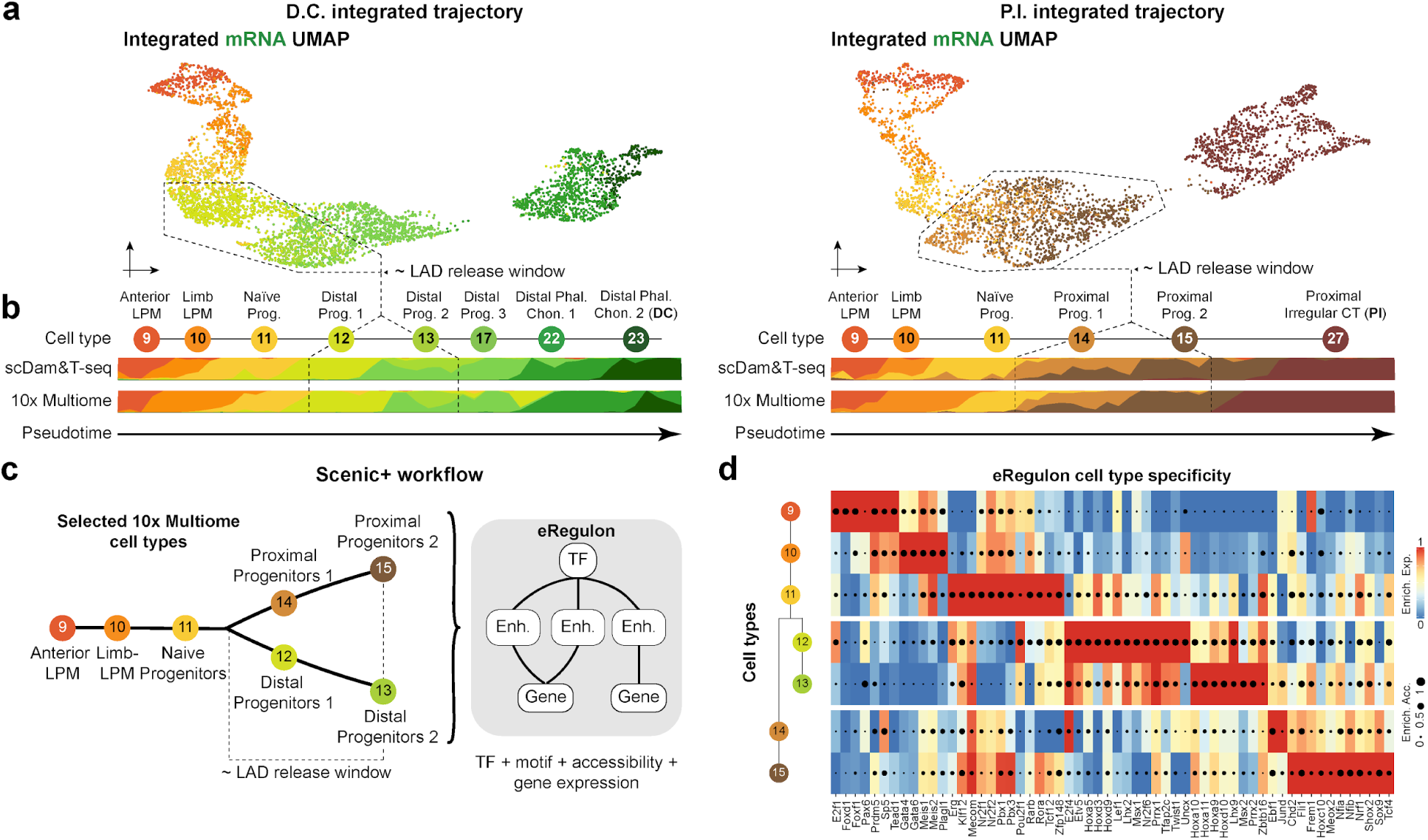
Integrated trajectory reconstruction and Scenic+ predictions of eRegulon activities and TF binding. **a**, UMAP visualisation of the integrated D.C. (n = 4544) and P.I. (n = 3572) trajectories based on the mRNA readout of the scDam&T-seq and 10x Multiome datasets. Cells are colored by cell type. **b**, Cell type distribution along pseudotime for the integrated trajectories shown in (a). **c**, Schematic of Scenic+ workflow. 7 LPM-derived cell types, up to the proximal and distal precursors where lamina-release is complete, were selected from the 10x Multiome dataset and used for eRegulon analysis. Each eRegulon is composed of a TF, the target regulatory elements it is predicted to bind, and the predicted genes it is inferred to control. **d**, Enrichment scores for Scenic+-identified eRegulons in indicated cell types. See also Figure 3.

**Extended Data Figure 9.**
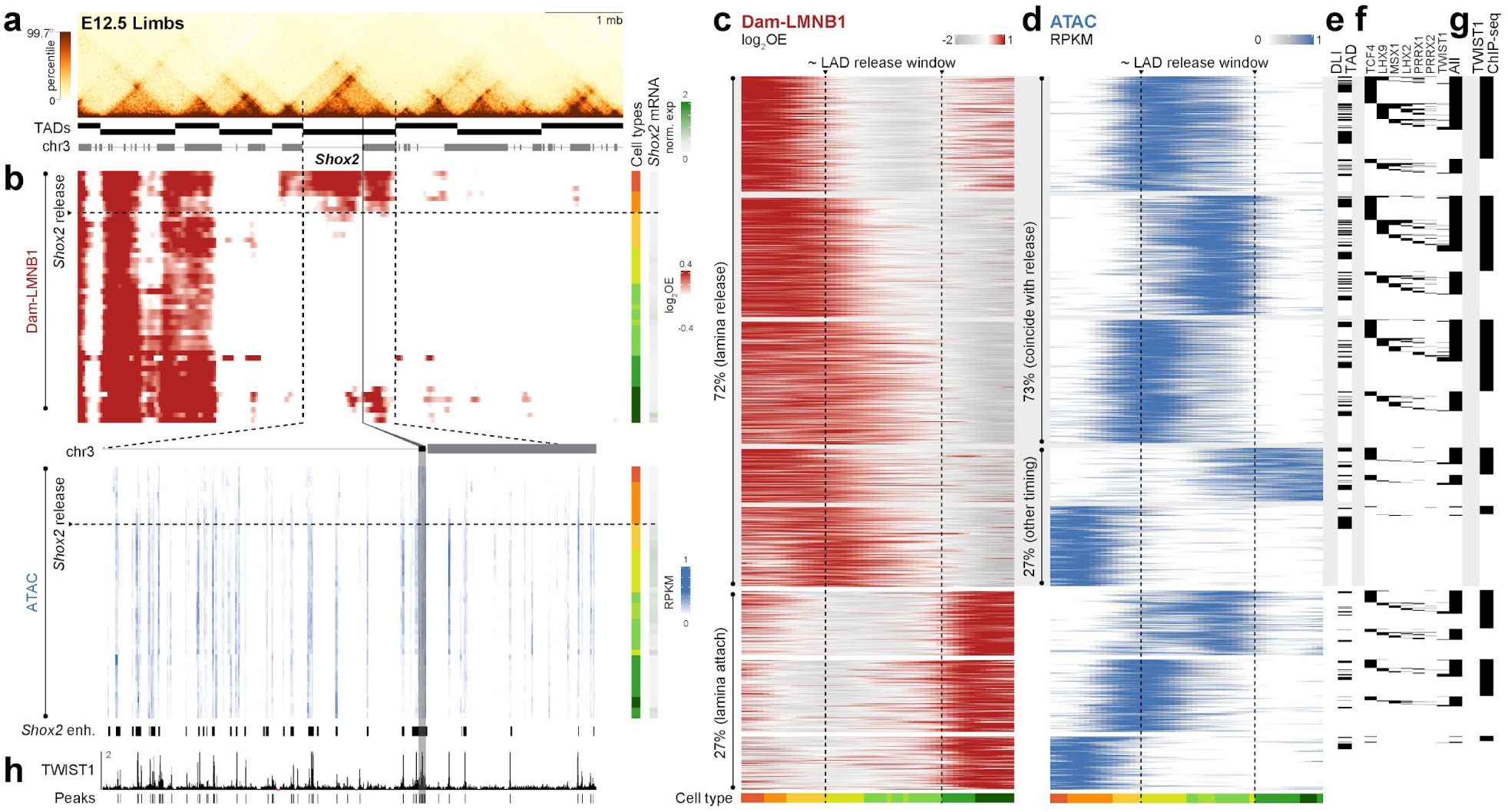
Additional LAD, chromatin accessibility, and expression dynamics in the D.C. trajectory. **a**, E12.5 limb Hi-C of the *Shox2* locus with called TADs (top) and gene track (bottom). **b**, Corresponding Dam-LMNB1 and ATAC signal (x axis) in single cells ordered along pseudotime (y axis) in the P.I. trajectory. *Shox2* mRNA signal per single cell is also shown (rightmost barplot). *Shox2* and its TAD are highlighted by the vertical gray bar and dotted lines, respectively. Lower single-cell ATAC heatmap is zoomed to the *Shox2* locus, with characterised *Shox2* enhancers shown below^46,47^. **c-d,** Heatmap of changing Dam-LMNB1 (c) and ATAC (d) signal for genomic bins that show both accessibility and lamina interaction dynamics (y axis) along D.C. trajectory pseudotime (x axis). Cell types are indicated below. **e-g,** Corresponding overlap of ATAC peaks with TADs of LAD-linked genes (e), predicted TF binding (f), or TWIST1 ChIP-seq peaks in E10.5 limbs^45^ (g). **h,** TWIST1 E10.5 forelimb ChIP-seq at the *Shox2* locus with called peaks below^45^. See also Figure 3.

**Extended Data Figure 10.**
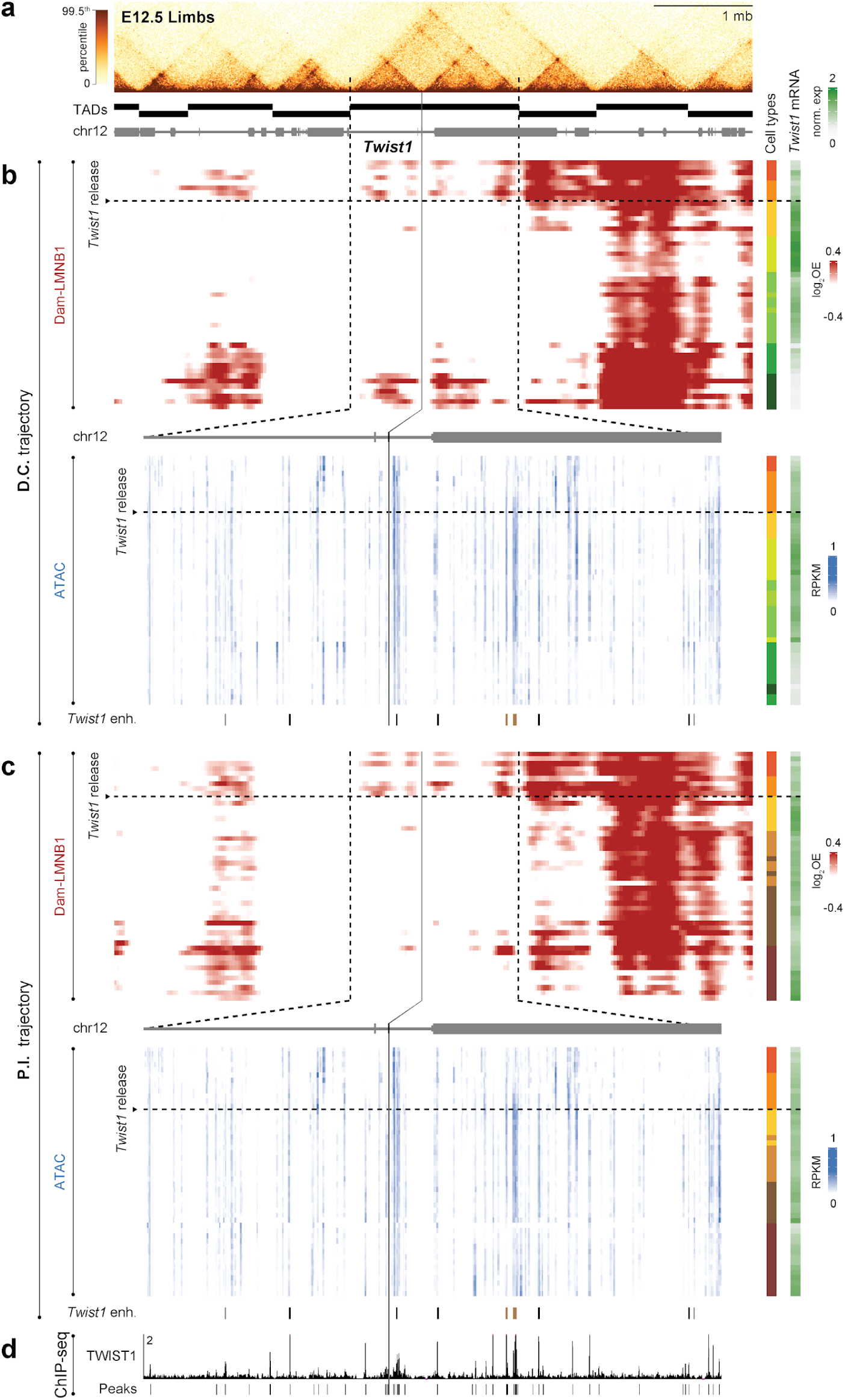
TWIST1 binds throughout the *Twist1* regulatory landscape during lamina-release. **a**, E12.5 limb Hi-C of the *Twist1* locus with called TADs. **b**, Corresponding pseudotime-ordered single cell profiles for Dam-LMNB1, ATAC and *Shox2* mRNA signal in the D.C. (left) or P.I. (right) trajectory. *Twist1* and its TAD are highlighted by the black and dotted vertical lines, respectively. Candidate (black bars) and validated (brown bars) Twist1 limb enhancers are also shown^55^. **c**, TWIST1 E10.5 forelimb ChIP-seq at the *Twist1* locus with called peaks below^45^. See also Figure 3.

**Extended Data Figure 11.**
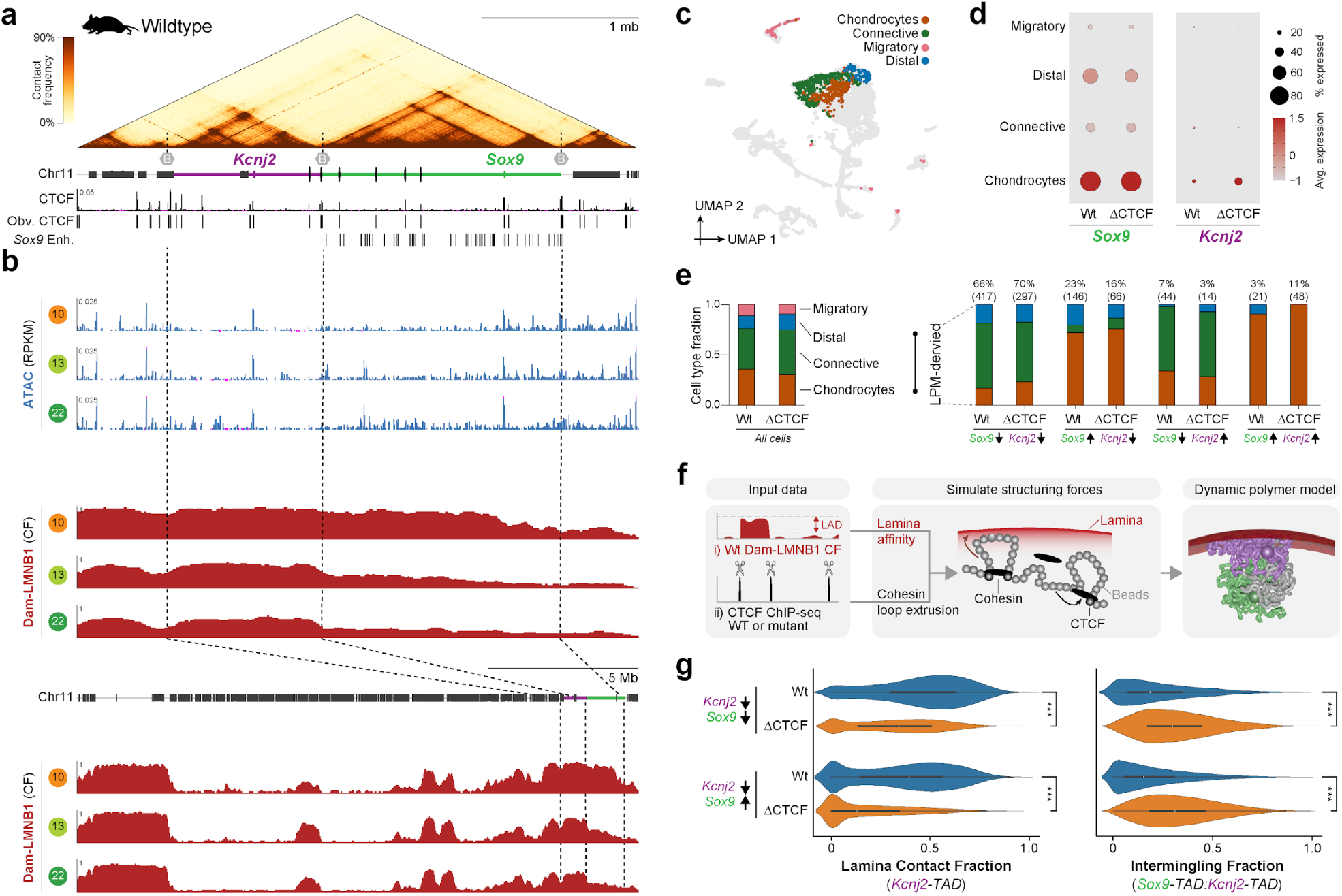
Expanded LAD and expression dynamics of the wild-type and ΔCTCF *Kcnj2/Sox9* locus. **a**, *Kcnj2/Sox9* cHi-C in E12.5 limbs with genes, CTCF ChIP-seq, CTCF peak-calling and Sox9 enhancers below^50,61^. **b**, Matching views of average scATAC-seq and Dam-LMNB1 signal for selected cell types from the D.C trajectory (upper and middle plots). Zoomed out view of scATAC-seq of the *Kcnj2/Sox9* locus, showing a broader region of chromosome 11 (lower plot). **c**, UMAP of wildtype and ΔCTCF E13.5 limb cells projected onto the primary scDam&T-seq dataset based on their mRNA readout. Colors and labels correspond to four major cell lineages at this stage of limb development. **d**, *Sox9* or *Kcnj2* expression across wildtype and ΔCTCF limb cell types. **e**, Broad cell lineage distribution of wildtype and ΔCTCF limb cells, including in LPM-derived cells grouped by low (↓) or high (↑) *Kcnj2*/*Sox9* expression levels. **f**, Schematic of beads-on-a-string polymer modelling approach. Lamina-association is modelled by assigning beads an affinity for a simulated lamina^29^ using wildtype Dam-LMNB1 contact frequency (CF) as input data. In parallel, TAD structures are simulated on the same polymer by randomly loading cohesin molecules to continuously extrude loops until blocked by CTCF barriers. Wildtype CTCF ChIP-seq^61^ defines the positions and frequency of CTCF binding, except in ΔCTCF simulations where selected CTCF sites are removed. See Methods for a detailed modelling description. **g**, Quantification of simulated structures with fraction of object in contact with the lamina (left) and object-object intermingling (right) shown. ***p < 0.001, from Welch’s t test comparisons between indicated samples. At least n = 8680 simulated structures per condition. See also Figure 4.

## SUPPLEMENTARY VIDEOS

**Video 1.** Representative polymer simulation of locus in wildtype limb cells with low Kcnj2 and high Sox9 expression.

**Video 2.** Representative polymer simulation of locus in ΔCTCF mutant limb cells with low Kcnj2 and high Sox9 expression.

## SUPPLEMENTARY TABLES

Table 1. Overview of collected samples for scDam&T

Table 2. Quality statistics for primary scDam&T dataset

Table 3. Literature used for annotation in this study

Table 4. Selected gene markers and literature used for annotating 60 scDam&T cell types

Table 5. DLI and DE status for all LAD-linked genes across five developmental trajectories

Table 6. Uncoupldness score for all LAD-linked genes across five developmental trajectories

Table 7. Overview of 10x Multiome runs performed for this study

Table 8. Overview of input material for 10x Multiome runs

Table 9. Quality statistics for 10x Multiome cells

Table 10. Basis of supervised 10x Multiome annotation based on scDam&T annotation and Seurat resolution 9

Table 11. Basis of supervised scDam&T annotation based on Seurat resolution 10

Table 12. Quality statistics for ΔCTCF mutants datasets

Table 13. List of ESC and mouse lines generated in this study

Table 14. Oligonucleotides used for genotyping in this study

## Notes

### Competing Interest Statement

The authors have declared no competing interest.

## REFERENCES

1. Arendt, D. et al. The origin and evolution of cell types. Nat. Rev. Genet. 17, 744–757 (2016).

2. Petit, F., Sears, K. E. & Ahituv, N. Limb development: a paradigm of gene regulation. Nat. Rev. Genet. 18, 245–258 (2017).

3. Zeller, R., López-Ríos, J. & Zuniga, A. Vertebrate limb bud development: moving towards integrative analysis of organogenesis. Nat. Rev. Genet. 10, 845–858 (2009).

4. McCarthy, R. L., Zhang, J. & Zaret, K. S. Diverse heterochromatin states restricting cell identity and reprogramming. Trends Biochem. Sci. 48, 513–526 (2023).

5. Schuettengruber, B., Bourbon, H.-M., Di Croce, L. & Cavalli, G. Genome Regulation by Polycomb and Trithorax: 70 Years and Counting. Cell 171, 34–57 (2017).

6. Bell, O., Burton, A., Dean, C., Gasser, S. M. & Torres-Padilla, M.-E. Heterochromatin definition and function. Nat. Rev. Mol. Cell Biol. 24, 691–694 (2023).

7. Padeken, J., Methot, S. P. & Gasser, S. M. Establishment of H3K9-methylated heterochromatin and its functions in tissue differentiation and maintenance. Nat. Rev. Mol. Cell Biol. 23, 623–640 (2022).

8. Bickmore, W. A. & van Steensel, B. Genome Architecture: Domain Organization of Interphase Chromosomes. Cell 152, 1270–1284 (2013).

9. Guelen, L. et al. Domain organization of human chromosomes revealed by mapping of nuclear lamina interactions. Nature 453, 948–951 (2008).

10. Ou, H. D. et al. ChromEMT: Visualizing 3D chromatin structure and compaction in interphase and mitotic cells. Science 357, eaag0025 (2017).

11. Poleshko, A. et al. Genome-Nuclear Lamina Interactions Regulate Cardiac Stem Cell Lineage Restriction. Cell 171, 573–587.e14 (2017).

12. Peric-Hupkes, D. et al. Molecular Maps of the Reorganization of Genome-Nuclear Lamina Interactions during Differentiation. Mol. Cell 38, 603–613 (2010).

13. Robson, M. I. et al. Tissue-Specific Gene Repositioning by Muscle Nuclear Membrane Proteins Enhances Repression of Critical Developmental Genes during Myogenesis. Mol. Cell 62, 834–847 (2016).

14. Robson, M. I. et al. Constrained release of lamina-associated enhancers and genes from the nuclear envelope during T-cell activation facilitates their association in chromosome compartments. Genome Res. 27, 1126–1138 (2017).

15. Shah, P. P. et al. An atlas of lamina-associated chromatin across twelve human cell types reveals an intermediate chromatin subtype. Genome Biol. 24, 16 (2023).

16. Shah, P. P. et al. Pathogenic *LMNA* variants disrupt cardiac lamina-chromatin interactions and de-repress alternative fate genes. Cell Stem Cell 28, 938–954.e9 (2021).

17. Marin, H. et al. The nuclear periphery confers repression on H3K9me2-marked genes and transposons to shape cell fate. 2024.07.08.602542 Preprint at 10.1101/2024.07.08.602542 (2024).

18. Lewis, R. et al. LBR and LAP2 mediate heterochromatin tethering to the nuclear periphery to preserve genome homeostasis. 2024.12.23.628302 Preprint at 10.1101/2024.12.23.628302 (2024).

19. Biferali, B. et al. Prdm16-mediated H3K9 methylation controls fibro-adipogenic progenitors identity during skeletal muscle repair. Sci. Adv. 7, eabd9371 (2021).

20. Finlan, L. E. et al. Recruitment to the Nuclear Periphery Can Alter Expression of Genes in Human Cells. PLOS Genet. 4, e1000039 (2008).

21. Reddy, K. L., Zullo, J. M., Bertolino, E. & Singh, H. Transcriptional repression mediated by repositioning of genes to the nuclear lamina. Nature 452, 243–247 (2008).

22. Brueckner, L. et al. Local rewiring of genome-nuclear lamina interactions by transcription. EMBO J. 39, e103159 (2020).

23. Therizols, P. et al. Chromatin decondensation is sufficient to alter nuclear organization in embryonic stem cells. Science 346, 1238–1242 (2014).

24. Tumbar, T. & Belmont, A. S. Interphase movements of a DNA chromosome region modulated by VP16 transcriptional activator. Nat. Cell Biol. 3, 134–139 (2001).

25. Kind, J. et al. Single-Cell Dynamics of Genome-Nuclear Lamina Interactions. Cell 153, 178– 192 (2013).

26. Long, H. K., Prescott, S. L. & Wysocka, J. Ever-Changing Landscapes: Transcriptional Enhancers in Development and Evolution. Cell 167, 1170–1187 (2016).

27. Robson, M. I., Ringel, A. R. & Mundlos, S. Regulatory Landscaping: How Enhancer-Promoter Communication Is Sculpted in 3D. Mol. Cell 74, 1110–1122 (2019).

28. Dekker, J. & Mirny, L. A. The chromosome folding problem and how cells solve it. Cell 187, 6424–6450 (2024).

29. Ringel, A. R. et al. Repression and 3D-restructuring resolves regulatory conflicts in evolutionarily rearranged genomes. Cell 185, 3689–3704.e21 (2022).

30. Gouhier, A. et al. Pioneer factor Pax7 initiates two-step cell-cycle-dependent chromatin opening. Nat. Struct. Mol. Biol. 31, 92–101 (2024).

31. Rooijers, K. et al. Simultaneous quantification of protein–DNA contacts and transcriptomes in single cells. Nat. Biotechnol. 37, 766–772 (2019).

32. Manzo, S. G., Dauban, L. & van Steensel, B. Lamina-associated domains: Tethers and looseners. Curr. Opin. Cell Biol. 74, 80–87 (2022).

33. Markman, S. et al. A single-cell census of mouse limb development identifies complex spatiotemporal dynamics of skeleton formation. Dev. Cell 58, 565–581.e4 (2023).

34. O’Rourke, M. P., Soo, K., Behringer, R. R., Hui, C.-C. & Tam, P. P. L. *Twist* Plays an Essential Role in FGF and SHH Signal Transduction during Mouse Limb Development. Dev. Biol. 248, 143–156 (2002).

35. Hui, C. & Joyner, A. L. A mouse model of Greig cephalo–polysyndactyly syndrome: the extra– toesJ mutation contains an intragenic deletion of the Gli3 gene. Nat. Genet. 3, 241–246 (1993).

36. Cobb, J., Dierich, A., Huss-Garcia, Y. & Duboule, D. A mouse model for human short-stature syndromes identifies Shox2 as an upstream regulator of Runx2 during long-bone development. Proc. Natl. Acad. Sci. 103, 4511–4515 (2006).

37. Singh, M. K. et al. The T-box transcription factor *Tbx15* is required for skeletal development. Mech. Dev. 122, 131–144 (2005).

38. ten Berge, D., Brouwer, A., Korving, J., Martin, J. F. & Meijlink, F. Prx1 and Prx2 in skeletogenesis: roles in the craniofacial region, inner ear and limbs. Development 125, 3831– 3842 (1998).

39. Michos, O. et al. Gremlin-mediated BMP antagonism induces the epithelial-mesenchymal feedback signaling controlling metanephric kidney and limb organogenesis. Development 131, 3401–3410 (2004).

40. Tzchori, I. et al. LIM homeobox transcription factors integrate signaling events that control three-dimensional limb patterning and growth. Dev. Camb. Engl. 136, 1375–1385 (2009).

41. Hoyt, P. R. et al. The *Evil* proto-oncogene is required at midgestation for neural, heart, and paraxial mesenchyme development. Mech. Dev. 65, 55–70 (1997).

42. Bi, W., Deng, J. M., Zhang, Z., Behringer, R. R. & de Crombrugghe, B. Sox9 is required for cartilage formation. Nat. Genet. 22, 85–89 (1999).

43. Otto, F., et al. *Cbfa1*, a Candidate Gene for Cleidocranial Dysplasia Syndrome, Is Essential for Osteoblast Differentiation and Bone Development. Cell 89, 765–771 (1997).

44. Schindler, M. et al. Comparative single-cell analyses reveal evolutionary repurposing of a conserved gene program in bat wing development. 2024.10.10.617585 Preprint at 10.1101/2024.10.10.617585 (2024).

45. Kim, S. et al. DNA-guided transcription factor cooperativity shapes face and limb mesenchyme. Cell 187, 692–711.e26 (2024).

46. Rouco, R. et al. Temporal constraints on enhancer usage shape the regulation of limb gene transcription. 2024.03.22.585864 Preprint at 10.1101/2024.03.22.585864 (2024).

47. Abassah-Oppong, S. et al. A gene desert required for regulatory control of pleiotropic Shox2 expression and embryonic survival. Nat. Commun. 15, 8793 (2024).

48. Bravo González-Blas, C., et al. SCENIC+: single-cell multiomic inference of enhancers and gene regulatory networks. Nat. Methods 20, 1355–1367 (2023).

49. Lallemand, Y. et al. Analysis of Msx1; Msx2 double mutants reveals multiple roles for Msx genes in limb development. Development 132, 3003–3014 (2005).

50. Despang, A. et al. Functional dissection of the Sox9–Kcnj2 locus identifies nonessential and instructive roles of TAD architecture. Nat. Genet. 51, 1263–1271 (2019).

51. Huang, X. et al. Single-cell, whole-embryo phenotyping of mammalian developmental disorders. Nature 623, 772–781 (2023).

52. Pal, M. et al. The establishment of nuclear organization in mouse embryos is orchestrated by multiple epigenetic pathways. Cell 0, (2025).

53. Isoda, T. et al. Non-coding Transcription Instructs Chromatin Folding and Compartmentalization to Dictate Enhancer-Promoter Communication and T Cell Fate. Cell 171, 103–119.e18 (2017).

54. Zakany, J. & Duboule, D. The role of *Hox* genes during vertebrate limb development. Curr. Opin. Genet. Dev. 17, 359–366 (2007).

55. Hirsch, N. et al. Unraveling the transcriptional regulation of TWIST1 in limb development. PLOS Genet. 14, e1007738 (2018).

56. Kind, J. et al. Genome-wide Maps of Nuclear Lamina Interactions in Single Human Cells. Cell 163, 134–147 (2015).

57. Sun, C., Zhao, Y., Guo, L., Qiu, J. & Peng, Q. The interplay between histone modifications and nuclear lamina in genome regulation. J. Genet. Genomics 52, 24–38 (2025).

58. Nicetto, D. et al. H3K9me3-heterochromatin loss at protein-coding genes enables developmental lineage specification. Science 363, 294–297 (2019).

59. See, K. et al. Lineage-specific reorganization of nuclear peripheral heterochromatin and H3K9me2 domains. Development 146, dev174078 (2019).

60. Wang, C. et al. Reprogramming of H3K9me3-dependent heterochromatin during mammalian embryo development. Nat. Cell Biol. 20, 620–631 (2018).

61. Andrey, G. et al. Characterization of hundreds of regulatory landscapes in developing limbs reveals two regimes of chromatin folding. Genome Res. 27, 223–233 (2017).

62. Beard, C., Hochedlinger, K., Plath, K., Wutz, A. & Jaenisch, R. Efficient method to generate single-copy transgenic mice by site-specific integration in embryonic stem cells. *genesis* **44**, 23–28 (2006).

63. Hayashi, S. & McMahon, A. P. Efficient Recombination in Diverse Tissues by a Tamoxifen-Inducible Form of Cre: A Tool for Temporally Regulated Gene Activation/Inactivation in the Mouse. Dev. Biol. 244, 305–318 (2002).

64. George, S. H. L. et al. Developmental and adult phenotyping directly from mutant embryonic stem cells. Proc. Natl. Acad. Sci. 104, 4455–4460 (2007).

65. Andrey, G. & Spielmann, M. CRISPR/Cas9 Genome Editing in Embryonic Stem Cells. in *Enhancer RNAs: Methods and Protocols* (ed. Ørom, U. A.) 221–234 (Springer, New York, NY, 2017). doi:10.1007/978-1-4939-4035-6_15.

66. Artus, J. & Hadjantonakis, A.-K. Generation of Chimeras by Aggregation of Embryonic Stem Cells with Diploid or Tetraploid Mouse Embryos. in *Transgenic Mouse Methods and Protocols* (eds. Hofker, M. H. & van Deursen, J.) 37–56 (Humana Press, Totowa, NJ, 2011). doi:10.1007/978-1-60761-974-1_3.

67. Markodimitraki, C. M. et al. Simultaneous quantification of protein–DNA interactions and transcriptomes in single cells with scDam&T-seq. Nat. Protoc. 15, 1922–1953 (2020).

68. Rang, F. J. et al. Single-cell profiling of transcriptome and histone modifications with EpiDamID. Mol. Cell 82, 1956–1970.e14 (2022).

69. Eberhart, A., Kimura, H., Leonhardt, H., Joffe, B. & Solovei, I. Reliable detection of epigenetic histone marks and nuclear proteins in tissue cryosections. Chromosome Res. 20, 849–858 (2012).

70. Durand, N. C. et al. Juicer Provides a One-Click System for Analyzing Loop-Resolution Hi-C Experiments. Cell Syst. 3, 95–98 (2016).

71. Li, H. & Durbin, R. Fast and accurate long-read alignment with Burrows–Wheeler transform. Bioinformatics 26, 589–595 (2010).

72. Shin, H. et al. TopDom: an efficient and deterministic method for identifying topological domains in genomes. Nucleic Acids Res. 44, e70–e70 (2016).

73. Langmead, B. & Salzberg, S. L. Fast gapped-read alignment with Bowtie 2. Nat. Methods 9, 357–359 (2012).

74. Kim, D., Paggi, J. M., Park, C., Bennett, C. & Salzberg, S. L. Graph-based genome alignment and genotyping with HISAT2 and HISAT-genotype. Nat. Biotechnol. 37, 907–915 (2019).

75. Hao, Y. et al. Dictionary learning for integrative, multimodal and scalable single-cell analysis. Nat. Biotechnol. 42, 293–304 (2024).

76. Grosswendt, S. et al. Epigenetic regulator function through mouse gastrulation. Nature 584, 102–108 (2020).

77. Pijuan-Sala, B. et al. A single-cell molecular map of mouse gastrulation and early organogenesis. Nature 566, 490–495 (2019).

78. Tao, H., Kawakami, Y., Hui, C. & Hopyan, S. The two domain hypothesis of limb prepattern and its relevance to congenital limb anomalies. *WIREs Dev*. Biol. 6, e270 (2017).

79. Zhao, W., Ji, X., Zhang, F., Li, L. & Ma, L. Embryonic Stem Cell Markers. Molecules 17, 6196– 6236 (2012).

80. La Manno, G. et al. Molecular architecture of the developing mouse brain. Nature 596, 92–96 (2021).

81. Zhao, L., Song, W. & Chen, Y.-G. Mesenchymal-epithelial interaction regulates gastrointestinal tract development in mouse embryos. Cell Rep. 40, 111053 (2022).

82. Jiang, S. et al. Single-cell chromatin accessibility and transcriptome atlas of mouse embryos. Cell Rep. 42, 112210 (2023).

83. Combes, A. N. et al. Single cell analysis of the developing mouse kidney provides deeper insight into marker gene expression and ligand-receptor crosstalk. Development 146, dev178673 (2019).

84. Mason, M. K. et al. Retinoic acid-independent expression of Meis2 during autopod patterning in the developing bat and mouse limb. EvoDevo 6, 6 (2015).

85. Mella, S., Soula, C., Morello, D., Crozatier, M. & Vincent, A. Expression patterns of the *coe*/*ebf* transcription factor genes during chicken and mouse limb development. Gene Expr. Patterns 4, 537–542 (2004).

86. Tahara, N., et al. *Gata6* restricts *Isl1* to the posterior of nascent hindlimb buds through *Isl1* cis-regulatory modules. Dev. Biol. 434, 74–83 (2018).

87. Yokoyama, S. et al. Analysis of transcription factors expressed at the anterior mouse limb bud. PLOS ONE 12, e0175673 (2017).

88. Mercader, N. et al. Ectopic *Meis1* expression in the mouse limb bud alters P-D patterning in a Pbx1-independent manner. Int. J. Dev. Biol. 53, 1483–1494 (2009).

89. Cao, J. et al. The single-cell transcriptional landscape of mammalian organogenesis. Nature 566, 496–502 (2019).

90. Buckingham, M. et al. The formation of skeletal muscle: from somite to limb. J. Anat. 202, 59– 68 (2003).

91. Negretti, N. M. et al. A single-cell atlas of mouse lung development. Development 148, (2021).

92. Feng, W. et al. Single-cell transcriptomic analysis identifies murine heart molecular features at embryonic and neonatal stages. Nat. Commun. 13, 7960 (2022).

93. Qiu, C. et al. A single-cell time-lapse of mouse prenatal development from gastrula to birth. Nature 626, 1084–1093 (2024).

94. Stuart, T., Srivastava, A., Madad, S., Lareau, C. A. & Satija, R. Single-cell chromatin state analysis with Signac. Nat. Methods 18, 1333–1341 (2021).

95. La Manno, G. et al. RNA velocity of single cells. Nature 560, 494–498 (2018).

96. Bergen, V., Lange, M., Peidli, S., Wolf, F. A. & Theis, F. J. Generalizing RNA velocity to transient cell states through dynamical modeling. Nat. Biotechnol. 38, 1408–1414 (2020).

97. Barbieri, M. et al. Complexity of chromatin folding is captured by the strings and binders switch model. Proc. Natl. Acad. Sci. 109, 16173–16178 (2012).

98. 98. Sanborn, A. L., et al. Chromatin extrusion explains key features of loop and domain formation in wild-type and engineered genomes. Proc. Natl. Acad. Sci. 112, E6456–E6465 (2015).

99. Fudenberg, G. et al. Formation of Chromosomal Domains by Loop Extrusion. Cell Rep. 15, 2038–2049 (2016).

100. Conte, M. et al. Loop-extrusion and polymer phase-separation can co-exist at the single-molecule level to shape chromatin folding. Nat. Commun. 13, 4070 (2022).

101. Chiariello, A. M., Annunziatella, C., Bianco, S., Esposito, A. & Nicodemi, M. Polymer physics of chromosome large-scale 3D organisation. Sci. Rep. 6, 29775 (2016).

102. Buckle, A., Brackley, C. A., Boyle, S., Marenduzzo, D. & Gilbert, N. Polymer Simulations of Heteromorphic Chromatin Predict the 3D Folding of Complex Genomic Loci. Mol. Cell 72, 786–797.e11 (2018).

103. Kremer, K. & Grest, G. S. Dynamics of entangled linear polymer melts: A molecular-dynamics simulation. J. Chem. Phys. 92, 5057–5086 (1990).

104. Allen, M. P. & Wilson, M. R. Computer simulation of liquid crystals. J. Comput. Aided Mol. Des. 3, 335–353 (1989).

105. Conte, M. et al. Loop-extrusion and polymer phase-separation can co-exist at the single-molecule level to shape chromatin folding. Nat. Commun. 13, 4070 (2022).

106. Chiariello, A. M. et al. Multiscale modelling of chromatin 4D organization in SARS-CoV-2 infected cells. Nat. Commun. 15, 4014 (2024).

107. Anderson, J. A., Glaser, J. & Glotzer, S. C. HOOMD-blue: A Python package for high-performance molecular dynamics and hard particle Monte Carlo simulations. Comput. Mater. Sci. 173, 109363 (2020).

